# Linking live-cell behavior to transcriptional responses across perturbations using dynamic caging

**DOI:** 10.64898/2026.05.05.723043

**Authors:** Brian Orcutt-Jahns, Raúl A. Reyes Hueros, Ana M. Meireles, Christian Cox, Luca Ghita, Jack Kamm, Cemre Celen, Yiming Yang, Eloise Zargari-Pariset, Nirvan Rouzbeh, Yiqi Zhou, Jie Wang, Shan Sabri, Takamasa Kudo, Pratiksha I. Thakore, David Richmond, Tommaso Biancalani, Hye-Jung Kim, Pier Federico Gherardini, Gary P. Schroth, Shannon J. Turley, Orit Rozenblatt-Rosen, Hector Corrada Bravo, Bo Li, Kathryn Geiger-Schuller

**Affiliations:** Genentech Research and Early Development, Genentech, Inc., South San Francisco, CA, USA; Cellanome, Inc., Foster City, CA, USA; Amgen, Inc., Thousand Oaks, CA, USA; Lila Sciences, Inc., San Francisco, CA, USA

## Abstract

Single-cell technologies, encompassing molecular, morphological, and functional assays, have emerged as cornerstones of modern biological research and discovery. However, current experimental methods often fail to explicitly link these ‘omic’ modalities, especially in live cells or longitudinally through time, impeding the study of multi-scale interactions and mechanisms of regulation. CellCage Enclosure (CCE) technology overcomes these limitations by dynamically compartmentalizing cells, allowing for scalable, live-cell, longitudinal exploration and simultaneous analysis of transcriptomic, proteomic, and morphological profiles. Using this novel technology, we generate previously inaccessible insights across various *in vitro* cellular systems under a diverse set of perturbations, including the discovery of morphological and proteomic features linked to immune suppressive gene set expression in human primary regulatory T cells (Tregs), as well as direct association of morphological and proteomic features with inflammatory gene modules in human colonic fibroblasts. We then develop a novel pooled CRISPR genetic screening technology using CCEs, PERTURB-LINK (PERTURBational LINKage of transcriptomics and imaging in single cells via enclosure-based screening) and apply this approach in murine bone marrow derived macrophages (BMDMs), enabling multiomic dissection of NF-κB pathway regulation in response to lipopolysaccharide (LPS) stimulation. Together, these findings demonstrate the broad impact that advancements in live-cell, paired multimodal technologies, especially upon perturbation, may offer in deepening our understanding of cellular biology.

## Introduction

The ability to probe biological systems at single-cell resolution has reached unprecedented scale across cell types, tissues, and organisms in *in vivo, in vitro,* and *in situ* settings. However, current single-cell technologies are unable to integrate live cell imaging to corresponding transcriptional states. Without the ability to link live-cell cellular behavior to transcription, previous technologies only provide a fragmented view of biological systems (Fig. 1A, right). Viewing biology from this perspective carries significant limitations; without deterministic linkage of readouts, linking cells across different imaging, expression, and functional modalities becomes impossible, making cross-modal mechanisms of regulation impractical to decipher, and obscuring our understanding of gene function, downstream signaling, cellular morphology, and causality.

**Fig. 1:**
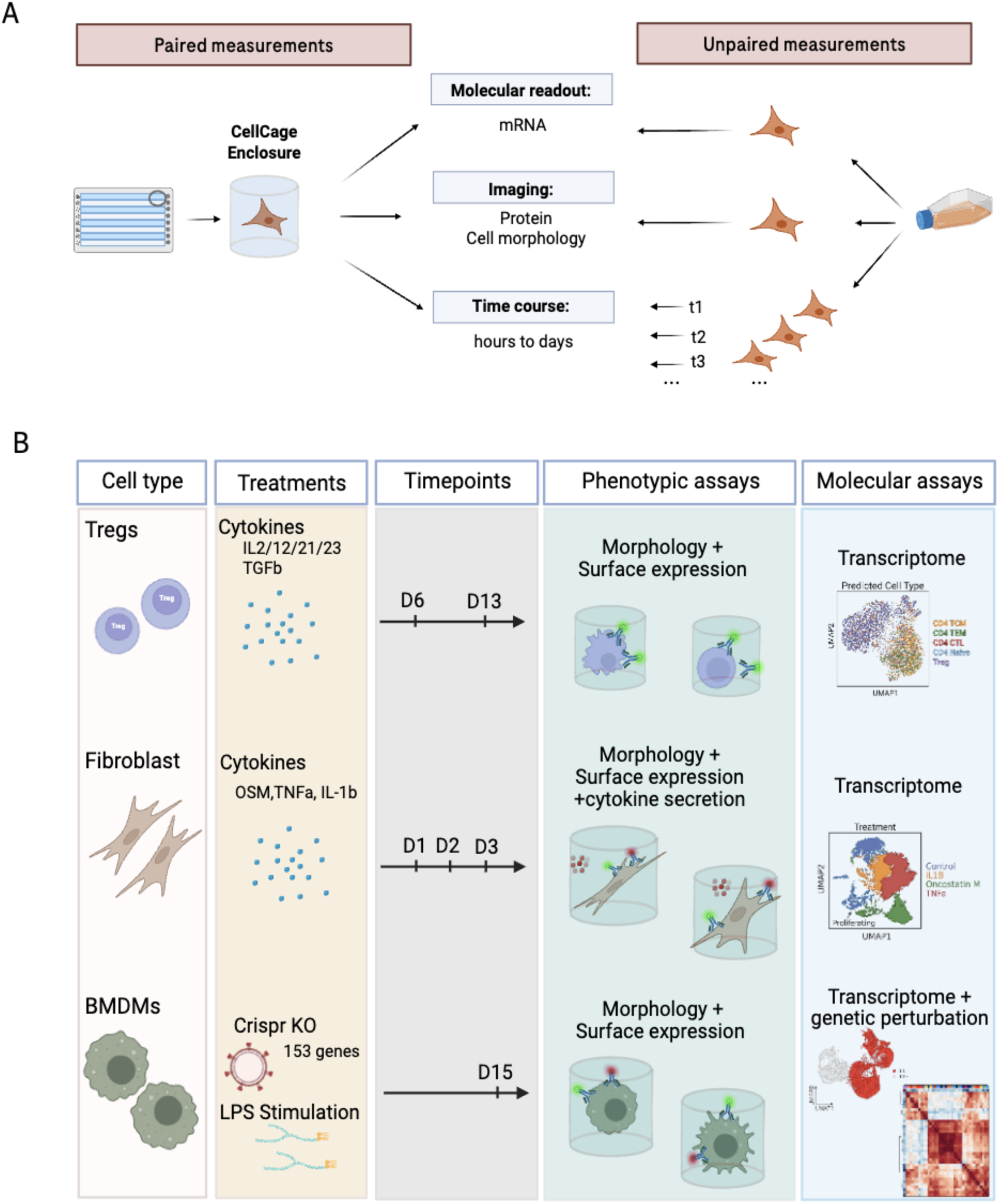
Unifying Omics and Imaging: a novel technology for comprehensive cellular analysis. **A,** Schematic depicting different levels of information obtained from traditional single cell RNA sequencing approaches, when used alone or in combinations (unpaired data, right side), vs how CCE allows integration of morphological, proteomic and transcriptomics data from the same single cell (paired data, left side). **B,** Summary schematic of the experiments performed, highlighting the various cell types used and different functional and molecular assays performed (Methods).

Recently developed technologies address specific aspects of pairing longitudinal imaging and transcriptomics in live cells, but face fundamental tradeoffs. Microwell- and precision barcoding-based methods allow for the linkage of imaging-readouts to transcriptomic profiles but only allow for single-timepoint imaging readouts^1–4^. Alternatively, semi-permeable capsule-based technologies generate both genomic and imaging readouts, but lack scalable deterministic linkage between images and transcriptomic profiles of single cells, limiting cross-modal insights to population-level inference^5,6^. Each of these approaches also lack dynamic, rule-based cell encapsulation, limiting utility for adherent cell or complex functional assays (eg. co-cultures, killing assays). Therefore, a need for a robust, high-throughput platform capable of true longitudinal paired profiling persists.

Here, we leverage the CellCage Enclosure (CCE) platform, described in Khurana et al. to overcome these limitations^7^. CCEs are hollow cylindrical compartments generated around cells via light-guided polymerization of a biocompatible hydrogel, enabling dynamic, rule-based capture and longitudinal imaging (Fig. 1A, left). This platform allows for direct, simultaneous, and high-throughput longitudinal acquisition of single-cell morphological, proteomic, and secretomic readouts via imaging of live suspended or adherent cells, paired to transcriptomic readouts, enabling previously inaccessible orthogonal profiling of *in vitro* systems. By leveraging CCEs we can more deeply explore the effects of various modes of perturbations, including small molecule, cytokine-based, and genetic perturbations, allowing deeper insights into causal gene function and cross-modal regulation (Fig. 1A, left).

In our study, we deployed CCEs in conjunction with cytokine perturbations to more fully characterize the response of suspended Tregs, to various cytokine treatments and identify links between morphological features and proteomic features, to Treg identity -gene programs. Second, we utilized CCEs to profile the response of adherent colonic fibroblasts to cytokine perturbations, where CCE-based profiling allowed us to identify both morphological features and protein markers which we directly link to upregulation of inflammatory gene programs, all without requiring the detachment of these cells at any stage. Lastly, we developed PERTURB-LINK (PERTURBational LINKage of transcriptomics and imaging in single cells via enclosure-based screening), a novel pooled CRISPR workflow that links genetic perturbations to their effects on cell morphology, protein expression and transcriptomic state. We applied PERTURB-LINK in bone marrow derived macrophages (BMDMs) for the simultaneous study of specific regulators of morphology, protein expression, and transcriptomic signatures driving the inflammatory response. Taken together, these applications demonstrate the profound value that simultaneous longitudinal imaging and paired transcriptional measurements offer in facilitating deeper biological insights from *in vitro* experimental settings under perturbation.

## Results

### Paired imaging and transcriptomic analysis of Tregs elucidates effects of cytokine stimulation, and reveals transcriptomically distinct T cell subsets distinguished by morphology and IL10 protein expression

To evaluate the utility of simultaneous multimodal analysis enabled by CCEs in the generation of integrated single-cell readout for suspension cells, we investigated the response of human regulatory T cells (Tregs) to various cytokine stimulations (Fig. 1B). Tregs, characterized by the expression of the master transcription factor Forkhead Box P3 (FOXP3), are central mediators of immune suppression, reducing inflammation through multifaceted pathways, including the secretion of immunosuppressive cytokines, making them attractive therapeutic targets and agents across various disease contexts^8–12^. Motivated by this, we perturbed Tregs under various cytokines and studied their response using the CCE platform. To profile the transcriptomic, proteomic, and morphological basis of Treg response, we activated human Tregs with T-cell TransAct, stimulated them with four distinct cocktails (conditions I-IV) comprised of various known Treg activators, IL-2 and TGF-β, as well as several factors, IL-12, IL-21, and IL-23 which have recently been reported to induce tissue-Treg states, and subsequently compartmentalized these cells in CCEs for multimodal analysis (Fig. 1B, 2A, Methods)^13–15^. CCE-based cell capture and profiling of morphological and transcriptomic data was performed on Tregs cultured for 6 days. After initial profiling, of the four conditions, two (conditions I and III) had remaining cells; to assess the CCE technology’s utility for evaluating functional cellular states as defined by protein expression, these remaining cells were re-stimulated for 7 days, and then stained for surface IL-10 protein using, followed by brightfield and fluorescent microscopy, and transcriptomic profiling. Rigorous quality control was conducted, and CCEs that were either empty or contained multiple cells were excluded; in total 21,922 high quality multiomic profiles were generated (Methods).

We first validated the quality of our data based on gene expression profiles.Transcriptomic UMAP embedding of all cells revealed two broad populations, primarily separated by FOXP3 expression (Fig. 2B). To determine whether these populations constituted functionally distinct subsets of Tregs, or true conventional T cell and classical Treg populations, we used CellTypist to transfer labels from the Hao et al. PBMC atlas^16,17^ (Methods). This approach confidently assigned the FOXP3-positive population as Tregs and the FOXP3-negative as predominantly T central memory (TCM) cells (Fig. 2B, Extended data Fig. 1A). Within our Treg population, cells clustered strongly by stimulation condition: Leiden clusters 1, 2, and 3 were chiefly composed of cells from Condition I, Conditions II/III, and Condition IV respectively (Fig. 2C). Cluster 1 represented cells at baseline and was not distinguished by unique gene expression patterns. Cluster 2 cells exhibited upregulation of *SATB1*, *CCL3*, and *ETS1*, suggesting an activated state, while Cluster 3 was distinguished by greater expression of *BATF3*, *CD70*, and various *MHC* genes, indicating that TGF-β induced a more suppressive state (Fig. 2D, Extended data Fig. 1B).

**Fig. 2:**
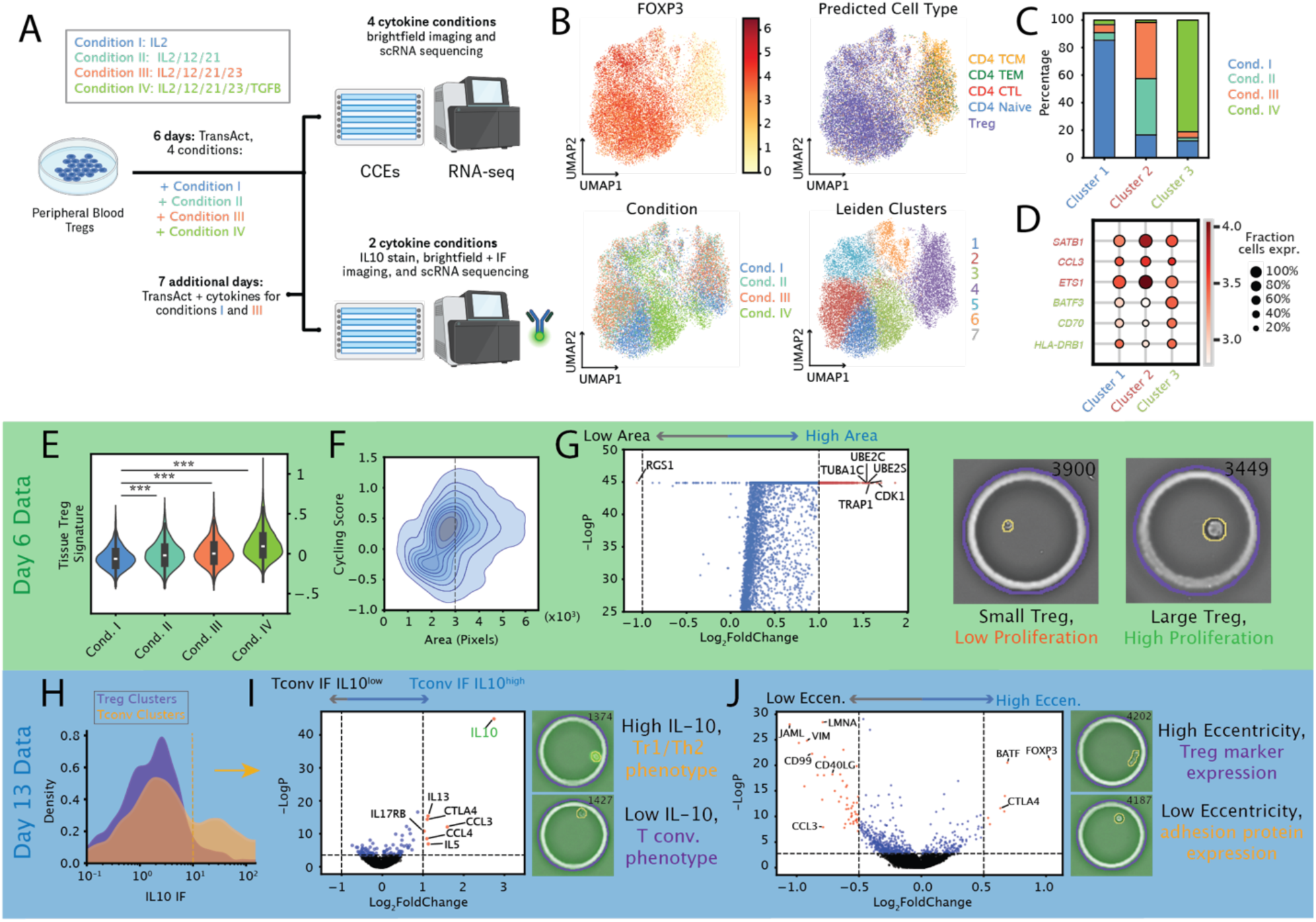
Paired imaging and transcriptomic profiling of human Tregs holistically characterizes response to cytokine stimulation. **A,** Schematic of Treg experimental design. Briefly, individual Tregs in CCEs were activated with T cell TransAct and stimulated with various cytokine cocktails for 6 days (Conditions I-IV) or 13 days (Conditions I+III). Tregs stimulated for 13 days were also stained for surface IL-10 protein levels. **B**, Uniform manifold approximation and projection (UMAP) of batch-corrected transcriptomic profiles of all 21,922 T cells from both 6 and 13 day time points colored by (left to right, top to bottom) FOXP3 expression, CellTypist predicted cell types from Hao et al. PBMC atlas (see methods)^17^, experimental stimulation condition, and Leiden cluster. **C**, Composition plot of experimental condition by Leiden cluster for clusters 1-3. **D**, Dot plot of Treg activation (red) and suppression (blue) markers for Leiden clusters 1-3. **E**, Violin plot of tissue Treg scores of cells profiled at day 6 in each experimental condition (see Methods). **F**, Kernel density estimate plot for cells profiled at day 6 of cell area, and cell cycling score (see Methods). Dotted line represents delineation between low area and high area cells in G. **G**, Volcano plot of differential expression (DE) analysis between large Tregs (cell area > 3000 pixels, i.e. diameter > 15.4 μm), and small Tregs (cell area < 3000 pixels). Red points: genes upregulated in large cells (right, FDR < 0.05, log2FC > 1), or genes upregulated in small cells (left, FDR < 0.05, log2FC < -1). Shown right are two brightfield images of cells typifying these classes. **H**, Kernel density estimate (KDE) plots of IL-10 immunofluorescence (IF) signal for cells profiled at day 13 in majority Treg clusters, and cells in majority Tconv. clusters (Extended data Fig. 1D). The dotted line represents delineation between low IL-10 and high IL-10 cells in I. **I**, Volcano plot of DE analysis between Tconv. cells with high IL10 IF signal (IL-10 IF > 10) and low IL-10 IF signal (IL-10 IF < 10). Red points: genes upregulated in IL-10 IF-high Tconvs. (right, FDR < 0.05, log2FC > 1). Shown right IF images of two cells typifying these classes. **J**, DE analysis between T cells profiled on day 13 with high eccentricity (eccentricity > 0.9) and cells with low eccentricity (eccentricity < 0.9). Red points: genes upregulated in highly eccentric cells (right, FDR < 0.05, log2FC > 0.5), or genes upregulated in low eccentricity cells (left, FDR < 0.05, log2FC < -0.5). Shown right are two IF images of cells typifying these classes. *\*Padj<0.05, **Padj<0.01, ***Padj<0.001*.

We next focused on the subset of cells which were profiled at day 6 (Extended data Fig. 1C). We scored our Treg clusters for the tissue Treg signature which was previously reported to be upregulated in conditions II and III compared to condition I (Methods)^15^. This signature was indeed upregulated in conditions II and III and, notably, was most strongly upregulated in condition IV (rank-biserial correlations of 0.16, 0.27, and 0.56 respectively), suggesting that TGF-β synergistically enhances tissue Treg phenotype when combined with IL-2, IL-12, IL-21 and IL-23 (Fig. 2E, Extended data Fig. 1C).

Having successfully performed unimodal analysis of our data, we next focused on harnessing the unique multimodality of our CCE-based approach to generate insights regarding relationships across transcriptomic, proteomic, and morphological readouts. We first explored linking single-cell morphological features, such as cell area, to interpretable gene programs. We observed a strong correlation between Treg cell size and a cell cycling gene signature score measured in the same single cell (Fig. 2F). Specifically, differential expression (DE) analysis comparing large Tregs (cell area > 3000 pixels, i.e. diameter > 15.4 μm) against all other Tregs revealed significantly increased expression of cell-cycle related genes, including *TUBA1C*, *TRAP1*, *UBE2C*, and *CDK1* (Fig. 2G). This highlights how the CCE technology enables the contextualization of cellular phenotypes, such as those defined by cell morphology, by explicitly linking them to transcriptomic features.

We characterized the cells profiled at 13 days and profiled surface IL-10 using the Miltenyi catch system (Methods, Extended data Fig. 1D). Given that signaling through IL-10 is a key mechanism by which Tregs exert their suppressive function, we were surprised to find that cells contained within our T conventional cell (Tconv) cluster contained more IL-10^+^ cells (IL-10 IF mean intensity > 10) than the Treg cluster (Fig. 2H). While this result was initially surprising, we also found that Tregs expressed little to no *IL10* in the Hao CD4 T cell atlas (Fig. 2H, Extended data Fig. 1E-F). To further understand this phenomenon, we performed DE analysis between IL-10^+^ Tconvs and other Tconvs (Fig. 2I). This analysis confirmed concordance between proteomic and transcriptomic readouts, revealing higher Type 1 regulatory T cell- (Tr1)- associated *IL10* and *CTLA4* mRNA alongside Th2-associated cytokines in FOXP3^-^ IL-10^+^ cells (*CCL3*, *CCL4*, *IL5*, *IL13*). FOXP3^-^ IL-10^+^ CD4^+^ T cells, or Tr1s, are well-documented in the literature as distinct suppressive populations, while Th2 cells are noted for their strong secretion of various pleiotropic cytokines^18,19^. Transcriptomic analysis of IL-10⁺ TCMs revealed enrichment of Tr1- and Th2-associated gene expression signatures compared to IL-10^-^ TCMs (Extended data Fig. 1G) along with elevated inferred activity of transcription factors EOMES and GATA3 (Extended data Fig. 1H), indicative of a hybrid or transitional cell state.

Finally, we further examined correlations between cell morphology and transcriptomic profiles on day 13. We consistently observed that Tregs exhibited higher eccentricity (less circularity) than TCMs (rank-biserial correlation 0.30) (Extended data Fig. 1I). This observation was confirmed via DE analysis comparing highly eccentric cells (eccentricity > 0.9) to other T cells, which revealed upregulation of canonical Treg markers in eccentric cells (*FOXP3*, *BATF*, *CTLA4*). Conversely, expression of several adhesive protein mRNAs (*VIM*, *JAML*, *CD99*) and T cell activation markers (*CCL3*) were significantly downregulated in highly eccentric cells (Fig. 2J). We also observed that mean eccentricity of condition I and III cells at day 6 was greater than that of condition I and III cells at day 13 (0.43 vs. 0.39, rank-biserial correlation 0.17), as was FOXP3 expression (log-normalized expression of 3.1 vs. 2.1, rank-biserial correlation 0.31), showing that the relationship between eccentricity and Treg marker expression was consistent across timepoints, and that corresponding alterations in morphology occurred alongside the expected gradual loss of Treg marker expression by Tregs over time in culture.

### Paired imaging and transcriptomic profiling of gut fibroblasts links proteomic and morphological features to inflammatory signatures

To evaluate the utility of our simultaneous multimodal analysis in adherent cells, we examined the temporal dynamics and cytokine response of human gut fibroblasts. Gut fibroblasts are key regulators of tissue repair and inflammation. Their central role in inflammatory bowel disease (IBD) signaling circuits has been increasingly studied, and therapies targeting the immune-fibroblast axis have advanced to clinical trials^20^.

To investigate transcriptomic, morphological, and proteomic aspects of gut fibroblast homeostasis and inflammatory responses, we profiled human Hs.675 colon fibroblasts using CCE-based multimodal readout (Fig. 3A). We conducted a time-series experiment, collecting high-quality morphological and transcriptomic readouts of 2,559 fibroblasts at time points of 0, 24, 48, and 72 hours (time course data and analysis also reported in companion manuscript, Khurana et al.)^7^. Concurrently, we performed a cytokine-response experiment, culturing fibroblasts with IL-1β, TNF, or Oncostatin M (OSM) for 72 hours before gathering transcriptomic and morphological data. We collected transcriptomic and morphological data from two biological replicates, and for one of the replicates we additionally included staining for surface markers (ICAM1, PDPN, PDGFRA) and profiling of IL-6 and CCL2 secretion using LEGENDplex cytokine capture beads. Again, exhaustive quality control excluded low-quality CCEs, as well as those that were empty or contained multiple cells (Methods).

**Fig. 3:**
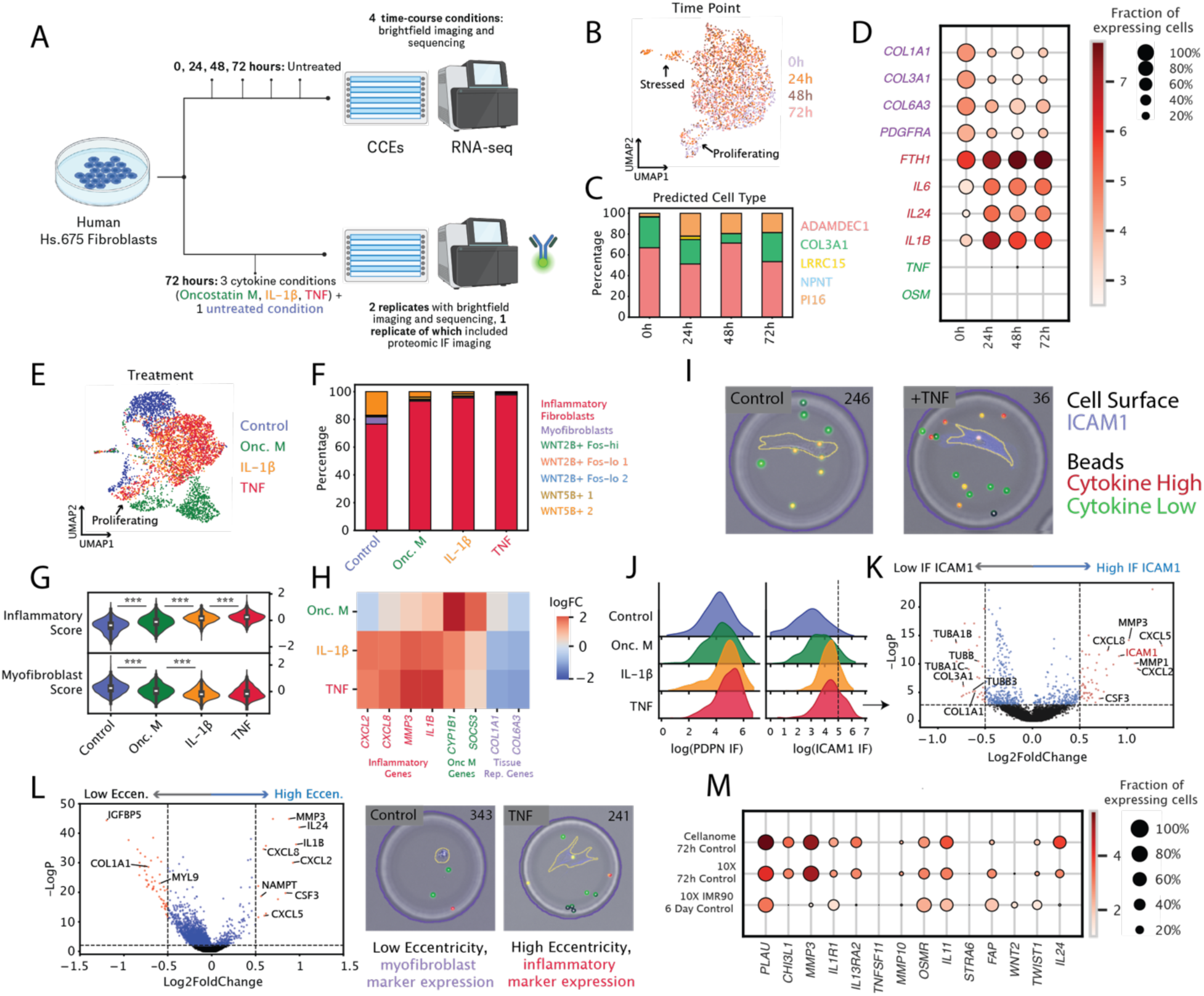
Paired imaging and transcriptomic profiling of colonic fibroblasts reveals proteomic and morphological markers of transcriptomically-defined inflammatory states. **A,** Schematic of human colonic fibroblast experimental design. Briefly, individual fibroblasts in CCEs were either left unstimulated, and profiled every 24 hours for 72 hours, or stimulated with one of three cytokines for 72 hours. One replicate of the 72 hour stimulated fibroblasts included IF-based imaging analysis of surface protein levels via fluorescent antibodies, and cytokine secretion via fluorescent beads. **B**, UMAP of batch corrected transcriptomic profiles of 2,559 time-course fibroblast cells colored by time point. **C**, Composition plot of the CellTypist-predicted cell types of fibroblasts profiled at each timepoint using the Buechler et al. fibroblast atlas^42^ (see Methods). **D**, Dotplot of expression of myofibroblast-related genes decreasing over time (purple), inflammation-related genes increasing over time (red), and cytokine-signaling-related genes (green) for fibroblasts profiled at various timepoints. **E**, UMAP of 4,598 fibroblasts in the stimulation experiment, colored by experimental condition. **F**, Composition plot of the CellTypist-predicted cell types of fibroblasts stimulated with various cytokines using the Smillie et al. fibroblast atlas^43^ (see Methods). **G**, Violin plot of inflammatory fibroblast and myofibroblast scores of fibroblasts treated with each cytokine stimulation (see Methods). **H**, Heatmap of cytokine-induced gene expression relative to untreated cells for fibroblasts in each experimental condition. **I,** IF images of untreated fibroblast and TNF-treated fibroblast. IF channels show surface markers (ICAM1, blue) and cytokine secretion bead activity. Yellow beads represent intermediate cytokine binding. LEGENDplex beads for IL6 and CCL2 are distinguished by bead size and fluorescence intensity (CCL2 smaller, more intense, IL6 larger, less intense). **J,** KDE plots of surface marker IF signal (PDPN, left, ICAM1, right) for cells from the stimulation experiment. Dotted line represents delineation between low ICAM1 and high ICAM1 cells in K. **K**, DE analysis between ICAM1-high fibroblasts (log(ICAM1 IF) > 6), and ICAM1-low fibroblasts (log(ICAM1 IF) < 6) across cytokine stimulation conditions. Red points: genes expressed more in ICAM1 IF-high fibroblasts (right, FDR < 0.05, log2FC > 0.5), or genes expressed more in ICAM1 IF-low fibroblasts (left, FDR < 0.05, log2FC < -0.5). **L**, Differential expression (DE) analysis between highly eccentric fibroblasts (eccentricity > 0.95), and less eccentric fibroblasts (eccentricity < 0.95) in the cytokine-response experiment. Red points: genes expressed in highly eccentric fibroblasts (right, FDR < 0.05, log2FC > 0.5), genes expressed more in more circular fibroblasts (left, FDR < 0.05, log2FC < -0.5). Shown right are two IF images of cells typifying the classes. **M**, Dot plot of inflammatory gene expression for control-group Hs.675 colonic fibroblasts profiled using CCE, untreated human Hs.675 colonic fibroblasts profiled using 10X, and untreated IMR90 normal, proliferating lung fibroblasts profiled using 10X from Palikyras et al.^44^. Hs.675 fibroblasts were cultured for 72 hours, while IMR90 fibroblasts were cultured for 144 hours. *\*Padj<0.05, **Padj<0.01, ***Padj<0.001*.

As in our study of Tregs, we first assessed whether our experimental data was suitable for biological discovery using unimodal analysis techniques. We first examined transcriptomic temporal dynamics (Fig. 3B, Extended data Fig. 2B-C), focusing on the time-series experiment. To validate that our fibroblasts phenotypically resembled colonic fibroblasts *in vivo*, we used CellTypist, to transfer labels from the Buechler et al. multi-tissue human fibroblast atlas^21^, and found that the majority of our fibroblasts were annotated as ADAMDEC1 gut fibroblasts, with smaller subsets identified as pan-tissue COL3A1 myofibroblasts and PI16 fibroblasts (Fig. 3C). We then assessed transcriptional changes in Hs.675 cells over time and observed a progressive downregulation of tissue-remodeling associated genes (*COL1A1*, *COL3A1*, *COL6A3*) and a concurrent upregulation of inflammatory markers (*IL6*, *IL24*) over time (Fig. 3D). This result was confirmed by pseudotime analysis, where inflammatory states and genes were inferred to become more prevalent over the 72-hour time course (Extended data Fig. 2E-F). Distinct upregulation of *IL1B* expression over time paired with consistently minimal *TNF* and *OSM* expression suggested that autocrine IL1β signaling may help explain this increasing inflammatory phenotype. Having characterized gut fibroblast homeostasis in this system, we next explored their response to inflammatory signals (Fig. 3A, bottom). Across the two biological replicates, we collected 4,598 high quality multiomic fibroblast profiles and observed that untreated fibroblasts clustered distinctly from cytokine-stimulated cells (Fig. 3E). We next dissected the fibroblast states further by using CellTypist with a gut-specific fibroblast atlas^22^ and found that a greater proportion of cytokine-stimulated cells were annotated as inflammatory fibroblasts, reflecting increased inflammatory polarization in these samples (Fig. 3F, Methods). We found that IL-1β and TNF induced a greater relative increase in inflammatory fibroblast scores compared to OSM (rank-biserial correlations of 0.65 and 0.74 vs. 0.32 respectively), as well as a greater decrease in myofibroblast scores (rank-biserial correlations of 0.51 and 0.45 vs. 0.25 respectively) (Fig. 3G-H). OSM also uniquely induced strong upregulation of genes associated with its signaling pathway, such as *SOCS3* and *CYP1B1* (Fig. 3H).

Having extensively profiled our data using traditional single cell analysis, we next sought to access the unique multimodal insights made possible by CCE-based analysis. We began by examining the relationship between proteomic and transcriptomic responses to inflammation. To study fibroblast secretion profiles, we used size and fluorescence encoded beads for CCL2 and IL-6 detection (LEGENDplex), demonstrating the utility of the CCE’s paired fluorescent imaging and dynamic cell and bead caging (Fig. 3I). Additionally, surface staining of ICAM1, PDPN, and PDGFRa proteins revealed that the expression level of each of these proteins correlated with the inflammatory gene signature across conditions, suggesting their utility as inflammatory markers (Fig. 3J, Extended data 2H). This observation was further evidenced by the analysis of ICAM1 protein-high (log(ICAM IF) > 6) versus ICAM1 protein-low cells, with ICAM1-high cells showing higher expression of *ICAM1* mRNA among other inflammatory genes (*CXCL5, CSF3, CXCL2, MMP1/3*), alongside downregulation of tissue remodeling genes (*COL1A1*, *COL3A1*) and various tubulins *(TUBB*, *TUBA1B, TUBA1C*), which may reflect proliferative or migratory states (Fig. 3K). Strikingly, we also found that ICAM1 protein expression correlated more significantly with inflammatory gene score than *ICAM1* mRNA expression (Spearman correlation of 0.41 vs. 0.17), highlighting how CCE’s proteomic readout may more effectively characterize functional states than transcriptomics alone.

Morphological changes also reflected transcriptional shifts in both the cytokine-response and time course cohorts. Highly eccentric cells (eccentricity > 0.95) exhibited increased inflammatory markers and decreased myofibroblast-related gene expression relative to fibroblasts at the lower tail of the eccentricity distribution, suggesting a link between cell morphology (adhesion, branching) and inflammatory gene expression (Fig. 3L, Extended data Fig. 2I-K). This association was also observed in our time-course data (Extended data Fig. 2I-K). Given the general upregulation of inflammatory genes over time, we investigated if this inflammatory progression was intrinsic to gut fibroblasts or a unique consequence of the cell’s interactions with the CCEs. We repeated the 72h-hours control condition experiment using Hs.675 fibroblasts on a separate 6-well tissue culture plate and profiled the cells using a 10x Genomics protocol (Methods). To further contextualize our findings, we also examined a public dataset of lung IMR90 fibroblasts cultured for 6 days prior to 10x profiling, which represented a healthy, non-pathogenic fibroblast population^23^. Hs.675 cells profiled via CCE showed high gene expression similarity to those profiled by 10x with respect to genes from the inflammatory signature. In addition, both were significantly more inflammatory than IMR90 cells, suggesting that spontaneous inflammation is an inherent feature of Hs.675 gut fibroblasts, and not an aberrant feature of CCE profiling (Fig. 3M, Extended data Fig. 2L-M).

### PERTURB-LINK uncovers functional architecture of the NF-κB pathway in macrophages

To demonstrate the unique capabilities of CCE-based paired-omics profiling in genetic perturbation screening, we developed PERTURB-LINK, a novel pooled CRISPR workflow that deterministically links genetic perturbations to their effects on cell morphology, protein expression, and transcriptomic state within the same single cells. We applied PERTURB-LINK to interrogate innate immunity in murine BMDMs, a cell type whose plasticity dictates both inflammatory and immunosuppressive responses to pathogens^24,25^. We used a curated 153-gene gRNA library targeting key regulators of immune function^26^, with a focus on the NF-κB signaling pathway and related inflammatory regulators. The library also included 10 olfactory receptor genes as cutting controls and 60 non-targeting controls (NTCs) (Fig. 4A, Supplementary Table 1). The NF-κB pathway is a central regulator of complex inflammatory responses and cell survival decisions of immune cells^27,28^, These processes involve mechanistic complexity that warrants simultaneous resolution of morphological, proteomic, and transcriptomic changes to establish causal relationships between perturbations and functional outcomes.. Following lentiviral transduction with our gRNA library, cells underwent puromycin selection and were enclosed within CCE. Subsequently, the cells were either stimulated with 100 ng/mL of lipopolysaccharide (LPS, a potent TLR4 agonist that robustly activates NF-κB signaling^29^) or mock-treated (LPS -) for 12 hours. We initially profiled 104,670 cells and simultaneously acquired single-cell transcriptomic, proteomic (CD40 and PD-L1 immunofluorescence), and detailed morphological features derived from brightfield microscopy images (Fig. 4A). After quality control, 64,093 cells were retained (61.23%), and 45,164 of these (70.46%) were confidently assigned to a valid gRNA (Methods).

**Fig. 4:**
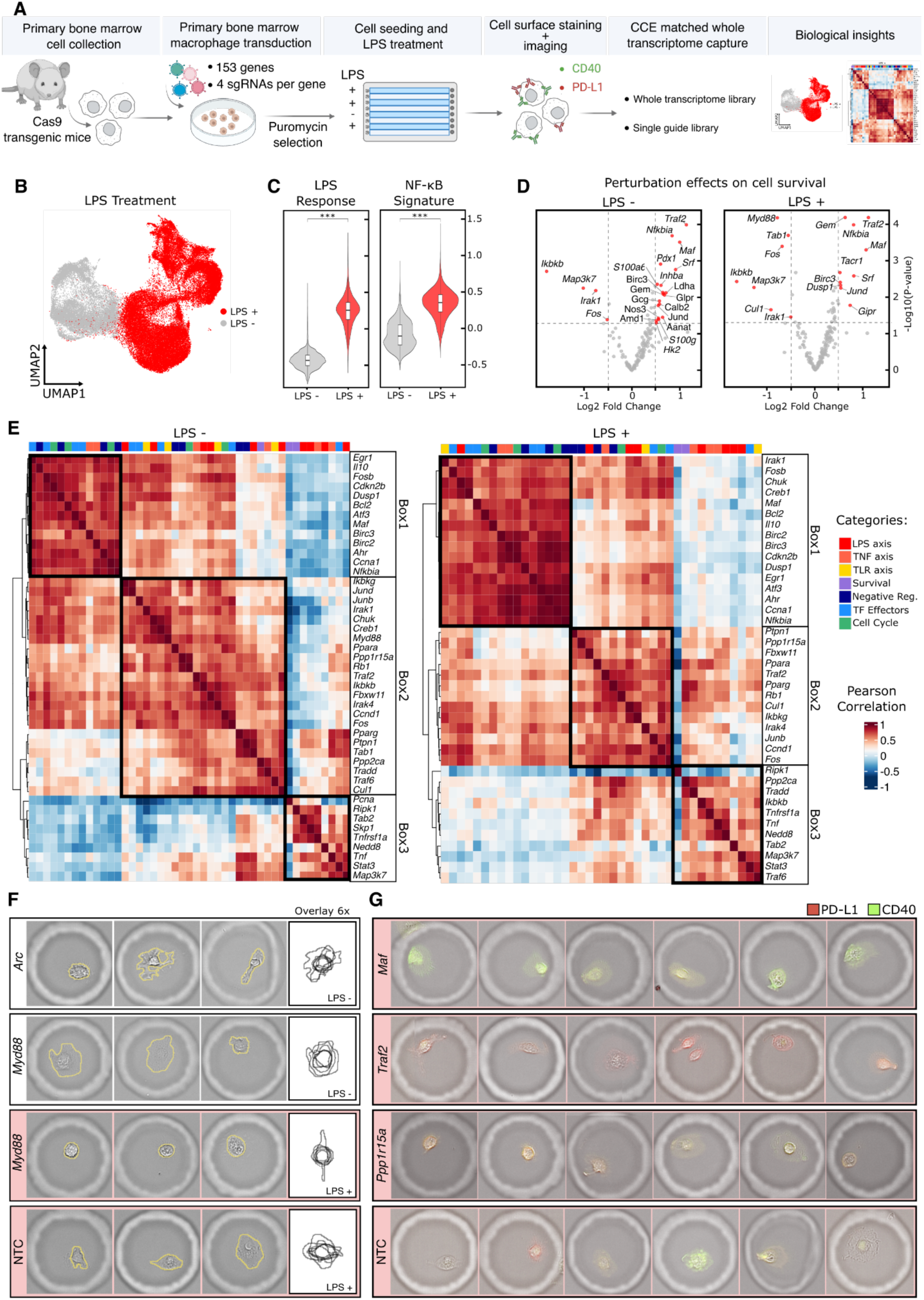
PERTURB-LINK in BMDMs dissects NF-κB pathway architecture. **A,** Schematic of the CRISPR knockout screen in murine BMDMs highlighting LPS stimulation conditions and paired-omics data collection (transcriptomics, proteomics, and morphology). **B,** UMAP embedding corrected by Harmony integration, highlighting mock-treated (LPS-, grey) and stimulated (LPS+, red) conditions. **C,** Violin plots showing the distributions of data-driven LPS response signatures from NTCs and literature-derived NF-κB pathway signature scores. Highly significant differences between LPS - and LPS+ conditions are indicated by significance (FDR < 0.05, log2FC > 0.5). **D,** Volcano plots of viability assay comparing gene perturbations to NTC in LPS- (left) and LPS+ (right) conditions. Red points indicate significant hits (FDR < 0.05, |log2FC| > 0.5). **E,** Heatmap illustrating Pearson correlations between gene perturbations in LPS– and LPS+ conditions, based on PCA of the collapsed beta-matrix from glmGamPoi modeling^50^. Hierarchical clustering annotations of sgRNA categories are labeled at the top of the heatmap. **F**, 3 representative brightfield images of BMDMs with DINOv2-identified morphological phenotypes, showing cells with altered boundaries and complexity following Myd88 and Arc knockout. The overlay is a graphical representation of cell masks generated from 6 cells. **G**, Immunofluorescence of CD40 (green) and PD-L1 (red) surface expression of select perturbations with significant proteomic phenotypes are highlighted. LPS – condition is represented in white and LPS + condition is represented in light salmon.

To validate PERTURB-LINK, we first examined the global transcriptomic response to LPS stimulation. A UMAP embedding of all batch-corrected single-cell profiles revealed a clear separation between LPS-treated and unstimulated populations (Fig. 4B). To specifically assess whether this separation was driven by known biology, we focused our analysis on NTC cells. LPS stimulation in NTCs induced a robust inflammatory activation, characterized by the upregulation of canonical NF-κB target genes such as *Tnf*, *Il1b*, and *Nfkbia*^30,31^. We then generated a data-driven “LPS response” signature by comparing stimulated versus unstimulated NTCs. Both the NF-κB and data-driven scores were significantly elevated in the LPS-treated cells, confirming a robust and canonical inflammatory response within our system (Fig. 4C; Extended data Fig. 3B-C). Interestingly, we observed baseline activation of NF-κB pathways even in unstimulated conditions (Extended data Fig. 3D), likely reflecting cell heterogeneity, cellular stress from lifting and seeding, lentiviral transduction and puromycin selection^32,33^.

As further validation of PERTURB-LINK, we assessed how genetic perturbations affected cell viability in both stimulated and unstimulated conditions (Fig. 4D). Viability was determined by quantifying the relative abundance of cells with each gRNA after 12 hours of culture, with a decrease in counts indicating a cytotoxic effect. Despite the relatively short 12-hour stimulation period, we identified clear patterns of stimulus-specific and shared gene dependencies. Knockout of *Myd88*, *Tab1*, and *Cul1* decreased macrophage viability under LPS stimulation but not in unstimulated conditions. This LPS-specific effect essentiality aligns with their known mechanistic roles: *Myd88* serves as the primary adaptor for TLR4-mediated NF-κB activation^29,34^, *Tab1* functions as a critical cofactor for TAK1 kinase activity^35,36^, and *Cul1* facilitates IκB degradation to enable NF-κB nuclear translocation^37^. Conversely, perturbation of core pathway components including *Ikbkb* (encoding IKKβ), *Map3k7* (encoding TAK1), *Irak1*, and the transcription factor *Fos* reduced cell survival regardless of stimulation status, highlighting their central roles across inflammatory/survival pathways^38,39^. The most significant perturbations that increased proliferation were *Traf2* and *Nfkbia*, which appear to enhance survival by disrupting the autocrine TNF-mediated apoptotic loop^40^. This is consistent with the known biology of activation-induced cell death, where strong inflammatory signaling can limit macrophage proliferation^24,25^.

As our final unimodal validation, we systematically mapped pathway relationships by assessing perturbation effects on gene expression using generalized linear models to generate a perturbation-by-gene beta matrix, where each value represents the estimated log-fold change of a gene’s expression resulting from a specific perturbation (see Methods for details). This matrix was then examined using correlation analysis. As expected, guides targeting expressed genes significantly regulated more genes and robustly downregulated the expression of their intended target gene compared to NTCs. As previously demonstrated in Perturb-seq datasets^41^, after aggregating guide-level effects to gene level, we found that 56 of the 153 targeted genes (∼36%) significantly regulated the expression of their intended target gene. (Extended data Fig.3B). The clustered perturbation-by-gene heatmap revealed clear groups aligned with known NF-κB pathway branches (Fig. 4E). For this visualization, we focused on 40 key perturbations targeting canonical inflammatory pathways (TNF, LPS, and TLR axes), downstream transcription factors, and related cellular processes like survival and cell cycle. Perturbations targeting core NF-κB signaling components clustered distinctly together (Fig. 4E, box 1), as did those affecting negative regulatory circuits and downstream transcriptional effectors (Fig. 4E, box 2). Perturbations linked to macrophage survival are closely associated with core NF-κB components, consistent with their critical roles identified in the viability analysis (e.g., *Ikbkb* and *Map3k7*, Fig. 4E, box 3). This integrative correlation analysis thus highlights the functional organization of the NF-κB signaling network, emphasizing the close interplay between pathway regulation, downstream effectors, and cellular survival mechanisms.

Having validated the screen’s performance using unimodal analyses, we next leveraged the CCE platform’s unique truly paired-omics capabilities. We first sought to identify novel links between genetic perturbations and cellular morphology, which CCEs enable by capturing both transcriptomic and imaging data from the same individual cell. To quantify broad morphological changes, we generated 1024-dimensional embeddings using the DINOv2 model^42^ from brightfield images and assessed perturbation-induced alterations using the energy distance statistic^72^. Our analysis resulted in 5 significant hits in LPS- condition and 25 significant hits in LPS+ condition (Extended data Fig. 4A-B). Among the significant hits, we highlighted top hits *Arc* in LPS- condition and *Myd88* in LPS+ condition as two examples (Extended data Fig. 4C). In unstimulated conditions, *Arc* knockout induced statistically significant morphological changes (Fig. 4F, permutation test, p-value<0.001). Although *Arc* has not been extensively characterized in macrophages, its established role in regulating actin-cytoskeleton dynamics in neurons and other myeloid cells like dendritic cells provides a plausible mechanistic basis for these alterations^44,45^. By contrast, *Myd88* knockout produced significant morphological changes in stimulated conditions, ranking among the top morphological hits, but not in the unstimulated (LPS -) conditions. Under LPS +, Myd88-deficient cells appeared more rounded with reduced spreading and structural complexity relative to controls (Fig. 4F; Extended data Fig. 4B). This pronounced stimulus-specific morphological phenotype aligns with Myd88’s essential role in TLR4-mediated NF-κB activation^60^, which is required for macrophage survival during LPS response. This pronounced stimulus-specific morphological phenotype aligns with Myd88’s essential role as the TLR4 adaptor that drives NF-κB-dependent anti-apoptotic gene expression ^53,60^, which protects macrophages from LPS-induced cell death.

We next extended this integrated analysis to our proteomic data, assessing how perturbation affected macrophage activation by quantifying two key surface proteins that fine-tune the immune response: the co-stimulatory molecule CD40, which amplifies inflammatory signaling, and the inhibitory checkpoint ligand PD-L1, which dampens it. Because both proteins were strongly induced by LPS, we focused our analysis on the stimulated condition, comparing each knockout to NTCs (Fig. 4G; Extended data Fig. 4C–E). In particular, we identified 9 and 5 hits for CD40 and PD-L1, respectively (p ≤ 0.05). Among the hits, we highlighted three top hits, *Maf*, *Traf2*, and *Ppp1r15a*, as examples (Fig. 4G, Extended data Fig. 4E). Several perturbations induced pronounced phenotypic changes in protein expression that complemented the transcriptomic data. *Maf* knockout, which increased viability in both stimuli, elevated CD40 under LPS, consistent with loss of c-Maf–dependent IL-10, an immunosuppressive cytokine that normally dampens the IFN-β–STAT1 signaling axis known to drive CD40 expression^46^. *Traf2* deletion robustly increased PD-L1, consistent with loss of TRAF2–cIAP restraint on non-canonical NF-κB and RIPK1-linked interferon signaling^40,47^. This pro-viability phenotype for both perturbations is consistent with the transcriptomic map (Fig. 4E), where Maf and Traf2 clustered separately from the cell-death associated group (box 3). *Ppp1r15a* knockout increased CD40 and PD-L1 with minimal impact on viability, which reflected in the single cell images (Fig. 4G, green and red fluorescence increases). This is consistent with loss of GADD34–PP1 restraint on TLR–TAK1/IKK and a prolonged eIF2α-dependent integrated stress response^48,49^. Together, these data recover canonical dependencies and nominate putative macrophage regulators. Notably *Arc*, which was identified through its morphological phenotype, and *Ppp1r15a,* a dual regulator of CD40 and PD-L1, suggesting previously underappreciated connections between cytoskeletal state, stress pathways, and checkpoint ligand expression which are uniquely revealed by CCE’s integrated readout.

## DISCUSSION

In this study, we harnessed CCE technology to generate rich paired-omics data under perturbation, which enabled novel insights across three diverse cellular contexts. Our single-cell-resolved transcriptomic, proteomic, and morphological readout first allowed us to thoroughly characterize the responses of human Tregs in suspension to tissue-cytokine factors. This application was of particular interest as, while current droplet-based transcriptomic and proteomic technologies are readily compatible with suspension cells, the simultaneous detection of gene expression and imaging features such as morphology and localization of protein expression in such cells remains challenging. There, we identified correlations between morphological features of live Tregs, such as eccentricity, with the induction of canonical Treg-associated gene expression. The proteomic readout was particularly valuable, enabling us to identify a subpopulation of potentially suppressive conventional T cells expressing IL-10 cytokines, a finding that, without the aid of immunofluorescence imaging, might have been misattributed to the sparsity often observed in transcriptomic data alone. Furthermore, in colonic fibroblasts, the platform’s single-cell resolution directly linked ICAM1 and PDPN protein expression to inflammatory gene module activity, offering new avenues for identifying these key cell types in inflammatory diseases. Live-cell imaging also allowed us to directly associate branching processes resulting in heightened fibroblast eccentricity with inflammatory gene expression, providing insight into the relationship between functional cellular processes and inflammatory gene expression. Studying adherent cells was particularly pertinent as many droplet or flow-based assays necessitate detachment of adherent cells, potentially introducing stress and aberrant signaling immediately preceding transcriptomic readout^51^, while CCEs allow for direct *in situ* lysis of adherent cell and in-cage primer-guided reverse transcription. Finally, we developed PERTURB-LINK, a novel CRISPR screening technology compatible with CCEs and applied it in BMDMs which enabled the deconvolution of complex gene functions, revealing genes whose impact was primarily on morphology (*Arc*) or on surface protein expression (*Ppp1r15a*), alongside core Nf-kB regulators (*Myd88*) whose essential roles manifested across all measured modalities. BMDMs and other myeloid cells, undergo known morphological changes in response to pathogens, and while previous work explored morphological changes and expression of select transcripts in the same single cells^52^, it was not previously possible to measure morphology and full transcriptome gene expression in the same single, live myeloid cell. While we were able to extensively validate the quality of each readout in isolation throughout our studies, the above findings, each of which would be inaccessible using single-omic approaches, demonstrate the deep biological value granted by expanding the breadth of data modalities examined in a focused experimental setting over time.

Single-cell, time-resolved multimodal analysis in *in vitro* settings as enabled by CCEs flexible cell capture, occupies a unique and complementary space in the current landscape of experimental tools at scientists’ disposal. As highlighted in the discussion of the advantages of microwell-, capsule-, and barcode-based technologies, many current platforms effectively address gaps in throughput, experimental flexibility, breadth of readout, and resolution depth; for example traditional spatial transcriptomic and proteomic technologies excel at profiling tissue organization at subcellular resolution, and several technologies have leveraged these platforms to generate paired transcriptional, proteomic, and morphological data^53–56^. CCEs build on this foundation by offering a platform specialized for dynamic stimulation, longitudinal observation, perturbation studies, and whole-transcriptome readouts. Imaging-based perturbation experiments allow for the linking of single-cell morphological states to genetic or chemical perturbations at scale^57^. CCEs extend these capabilities by deterministically linking full transcriptional and morphological states of single cells under perturbation, potentially helping contextualize the morphological responses observed using imaging-based techniques. Live-cell RNA resampling technologies address the need for longitudinal transcriptomic measurements; CCEs advance this approach by linking longitudinal live-cell imaging to transcriptional states at high throughput^58,59^. Other approaches have harnessed DNA nanoball technology to enable single-cell-resolved multiomic readouts^60^. CCEs add a new dimension to this technology by allowing for dynamic rule-based cell encapsulation, enabling the study of secretomics via immunofluorescent analysis of fluorescent beads and multi-cellular functional assays. As demonstrated by this study, CCEs represent a unique tool by blending temporally resolved analysis of live-cell morphology, gene, and protein expression which confers unique advantages in the terms of the breadth and depth at which myriad *in vitro* systems can be studied.

While this study has demonstrated many of the advantages of dynamic CCE-enabled multimodal readout, many opportunities to expand its utility, both experimentally and computationally, remain. Our current applications have focused on encapsulation of single cells. However, future studies could aim to capture multiple cells, enabling rigorously controlled studies of cell-cell interactions and thus more sophisticated functional assays with integrated morphological, proteomic, and genetic perturbation readouts, especially building off of the perturbation studies enabled by PERTURB-LINK. Current technologies largely interrogate cell-intrinsic gene regulation, whereas our approach enables scalable analysis of how genetic perturbations in one cell influence morphology, function, and transcriptional state of neighboring cells, providing a framework to study complex multicellular interactions such as Treg-mediated immunoregulation impacting cytotoxic T cell killing and fibroblast-macrophage communication. The application of transcriptomic deconvolution algorithms, aided by ground-truth cell-type assignments, may also enable meaningful transcriptomic readouts in these multi-cell contexts^61–63^. Computationally, further integration of imaging-based and transcriptomic readouts via weighted-neighbor, deep-learning, or dimensionality-reduction-based methods could potentially enhance the resolution of cellular identity and uncover novel cell types and states which are only possible to define in multimodal space^16,64–66^. Additionally, the ability to perform cell fixation and subsequent staining combined with high-content microscopy could facilitate richer morphometric analysis^57^.

In summary, the use of CCEs for paired imaging and omics readouts in single cells represents a significant step forward in the ongoing pursuit of richer, higher-content data. The technology’s blend of integrated omics readouts which can be harnessed flexibly across experimental settings, especially including this development of pooled perturbational studies, is representative of the innovation needed to enable the next generation of biological discovery, as exemplified by this paper’s findings.

## ACKNOWLEDGEMENTS

We would like to thank Jake Yeung for his advice on analyzing the perturbation screen data. We thank Aviv Regev for her support, insights and helpful discussions. We further owe thanks to the End-to-End Data team at Genentech for making this study possible. We additionally thank Zora Modrusan, Kevin Huang, Josh Kaminker, and Jason Rock for their insightful feedback on the manuscript. Schematic figures were created using BioRender.com.

## AUTHOR CONTRIBUTIONS

A.M.M., K.G.S., S.J.T., C.C., L.G., and H.J.K. conceived the study. J.W. performed the Treg characterization experiments. N.R., and C.C. performed the fibroblast characterization experiment. A.M.M., E.Z.P., and Y.Z. performed the BMDM perturbation screen, and T.K. provided further experimental support. B.O.J. performed the analysis of the Treg and fibroblast studies under the guidance of B.L.. R.A.R.H. performed analysis of the perturbation screen under the guidance of K.G.S. and H.C.B.. J.K, Y.Y., S.S. and P.F.G. provided computational support for the study. B.O.J., A.M.M., R.A.R.H., B.L., and K.G.S. wrote the manuscript with, input from all authors. B.L., H.C.B, K.G.S., P.I.T., S.J.T., D.R., T.B., O.R.R, G.P.S. supervised the work.

## CODE AND DATA AVAILABILITY

All data and analysis contained in this paper will be made public at the time of this report’s publication and will be deposited in a public repository such as Zenodo.

## METHODS

### EXPERIMENTAL METHODS

#### Generation and Cell culture within CellCage™ Enclosures (CCEs)

The CCE workflow starts with cells in suspension, which are mixed with a biocompatible hydrogel precursor mix and loaded into an 8-lane flow cell. Following an imaging scan and leveraging computer vision within the instrument, the optimal placement of compartments containing cells and barcoded capture oligos is determined. Polymerization is then induced by projecting 405 nm light to form the desired enclosure pattern. The enclosed cells can be cultured and imaged for several days before cell lysis and transcriptome capture occurs (see companion manuscript Khurana et al.)^7^.

#### Treg treatment and imaging

Human Peripheral Blood CD4^+^CD25^+^CD127lowFOXP3^+^ Treg cells were purchased from Stemcell (#200-0120). 200,000 cells per well were activated with anti-human T cell TransAct (Miltenyi Biotec, 130-111-160) and cultured in TexMACs medium (Miltenyi Biotec, 130-097-196) under four different conditions: (1) IL-2 only (PeproTech, #200-02-10UG, 500 U/mL); (2) IL-2 (500 U/mL), IL-12 (PeproTech, # 200-12P80H-10UG, 50 ng/mL), and IL-21 (PeproTech, # 200-21-10UG, 50 ng/mL); (3) IL-2 (500 U/mL), IL-12 (50 ng/mL), IL-21 (50 ng/mL), and IL-23 (PeproTech, # 200-23-10UG, 50 ng/mL); and (4) IL-2 (500 U/mL), IL-12 (50 ng/mL), IL-21 (50 ng/mL), IL-23 (50 ng/mL), and TGF-β (PeproTech, # 100-21-10UG, 50 ng/mL). After 3 days of culture, TransAct was removed, and cytokines were re-supplemented for an additional 3 days of incubation. On day 6, Tregs from all four cytokine conditions were resuspended in Cellanome hydrogel precursors and immediately processed for mRNA capture following CCE formation.

In parallel, the remaining Tregs from condition 1 were re-activated on day 6 using TransAct in the presence of IL-2 (500 U/mL), and those from condition 3 were re-activated with TransAct in the presence of IL-2 (500 U/mL), IL-12, IL-21, and IL-23 (each at 50 ng/mL). On day 13, IL-10 secretion was assessed using the Miltenyi IL-10 Secretion Assay (Cat# 130-090-434). Tregs were labeled with IL-10 catch reagent according to the manufacturer’s instructions, loaded onto flowcell, and stimulated with TransAct to induce cytokine secretion. After 20 hours of incubation at 37 °C, PE-conjugated IL-10 detection antibody (1:200 dilution) was added. Imaging was performed using the Cellanome R3200 instrument, followed by cell lysis and mRNA capture for downstream analysis.

#### Fibroblast treatment and imaging

Hs 675.T human colon fibroblasts (ATCC; CRL-7400) were cultured in Dulbecco’s Modified Eagle Medium (DMEM; ATCC, 30-2002) supplemented with 10% fetal bovine serum (FBS; ATCC, 30-2020), 1% penicillin-streptomycin (ATCC, 30-2300), 5 μg/mL insulin (Lonza Biosciences, BE02-033E20), and 10 ng/mL FGF-2 (R&D Systems, BT-FGFBHS-010). Cells were maintained at 37°C in a humidified incubator containing 5% CO₂.

On the day of the experiment, fibroblasts were detached using Trypsin-EDTA (Thermo Fisher Scientific, 25200072) and resuspended in cell media containing Cellanome hydrogel precursors mixed with IL-6 and CCL2 LEGENDplex cytokine capture beads (BioLegend; 740800 and 740200, respectively). These beads are size and fluorescent dye encoded allowing for easy demultiplexing of captured signal. The cell-bead mixture was loaded onto a Cellanome 3T genomic flow cell and transferred to the Cellanome R3200 instrument for CCE formation. Flow cells were incubated at 37°C with 5% CO₂ for 72 hours, with daily media exchanges.

Following incubation for multimodal analysis, cells were stained with fluorescently conjugated antibodies against IL-6 (BioLegend, 504504), CCL2 (BioLegend, 502604), podoplanin (PDPN; BD Biosciences, 753243), PDGFRα (BD Biosciences, 562798), and ICAM-1 (SC Biotechnology, sc-18908 FITC). Imaging was performed using the Cellanome R3200 instrument, then cells were lysed and processed for mRNA capture. In parallel, cells cultured on separate cell culture plates, were collected and lysed for mRNA capture using a Single Cell 3’ Reagent Kit (v3) on a Chromium (10X genomics) instrument.

#### BMDM Cell Culture

Bone marrow was collected from tibias and femurs of 8- to 14-week-old Cas9–enhanced green fluorescent protein (eGFP) transgenic mice. Following red blood cell lysis using ACK lysis buffer (Gibco, A1049201), 10 million cells were plated in BMDM medium (DMEM high glucose, 10% Tet-Negative heat-inactivated FBS, 1× GlutaMAX, 100 U/mL penicillin–streptomycin and 50 ng/mL recombinant mouse macrophage colony-stimulating factor (Genentech media facility) at a density of 1 million cells per ml in 10cm non tissue culture treated dishes (ref 351146, Corning). On day 3 after plating, plates were washed once with PBS and fresh media and pooled lentivirus supernatant was added to cells overnight (Supplementary Table 1). Transduced cells were selected with 5ug/ml puromycin from day 8 to day 11. For loading onto the instrument, cells were washed once with PBS and incubated in 5mL of Accutase (ref 7920, Stem Cell Technologies) at 4C for 10 min. Cells were collected, spun down and resuspended to a density of 1.5e6 cells/mL in media with 100ng/mL of mouse macrophage colony stimulating factor.

#### LPS Treatment and immunostaining for BMDM PERTURB-LINK

Cells were stimulated with 100ng/mL LPS (ALX-581-010, Enzo Life Sciences 1mg/mL) for 12hrs. After LPS stimulation cells were washed with PBS and blocked with 2.5ug/mL TruStain FcX plus (cat nr 156604) in Staining Buffer (1x PBS, 2% BSA, 0.01% Tween) for 10 min at RT. 200 μL of Staining buffer with a 1:25 and 1:25 dilution of PE-conjugated anti Cd40 antibody (cat nr 124610) and APC conjugated anti-PD-L1 antibody (cat nr 124312) was loaded onto the lane. Cells were incubated for 30 min at room temperature and washed 3 times before imaging.

### COMPUTATIONAL METHODS

#### Transcriptomic preprocessing pipeline

Processing of sequencing data was performed as described in our technology companion paper, Khurana et al.^7^. Briefly, STARSolo^67^ was used to generate barcodes by gene count-matrices. Counts are then aggregated per CCE by summing transcript counts. CCEs are then filtered based on CCE-barcode overlap to ensure efficient capture of mRNA. Upon aggregation, quality control was performed using standard workflows combining transcriptomic- and imagining-based quality controls. Briefly, in Treg experiments cages with >750 MBCs and <15% mitochondrial reads were kept, and in fibroblasts, cages with >1000 MBCs and <20% mitochondrial counts were kept. Quality control steps in BMDMs are described below. After CCE filtering, preprocessing, and CCE-level aggregation was performed using SeqAnalysis, an R package specifically developed for analysis of data from CCE technology.

#### Image analysis and DINOv2 embedding

Imaging analysis was performed as described in our technology companion paper, Khurana et al.^7^. Briefly, YOLOv5^68^ was trained on 15,000 images in order to generate object bounding boxes during CCE formation, and Mask R-CNN^69^ was trained and used to generate detailed object masks quantifying imaging features. Segmentation of objects was performed on brightfield images. Immunofluorescence quantification was calculated as the mean pixel intensity per object, and background subtraction was performed. The DINOv2^42^ model was used to generate cell-level morphology embeddings.

### Analysis Pipeline

#### Analyses of Treg and Fibroblast and CCE-Aggregated Datasets in Python

Treg and fibroblast datasets were analyzed using a combination of custom Python code and Pegasus v1.10.2^70^. First, CCEs with exactly one cell were isolated, and CCEs with 0, >1, or unknown numbers of cells were filtered out. Second, raw counts were normalized to CPK100K (counts per 100,000) at the single CCE level, and log-transformed (log(CP100K+1). Thirdly, the top 2,000 highly variable genes (HVGs) were identified, and 50 principal components were used for downstream analysis. In cases where batch correction was performed, the effects of the number of genes was regressed out, and then Harmony-Pytorch^71^, a Pytorch implementation of the Harmony batch correction algorithm^72^ was applied with default parameters and GPU mode. The batch keys used for each integration were as follows: Fig. 2A: Treg cohort (6 days treatment, 13 days treatment), Fig 3B: Fibroblast time point (0h, 24h, 48h, 72h), Fig. 3C: experimental replicate (Fibroblast treatment replicate 1, replicate 2). Using either the uncorrected PCs, or the Harmony-corrected PCs, a k-nearest neighbor graph was then built, and utilized to construct UMAPs^71^ for single cells, as well as for cluster detection using the Leiden community detection algorithm^73,74^. Cell type annotations were predicted using CellTypist as described below. Gene program signature scores were calculated using Pegasus and the exact gene list for each signature is listed in extended data table 1. Differential expression analysis between cellular populations was performed using the Mann-Whitney-U test implemented in Pegasus. The false discovery rate for genes was controlled at 5% using the Benjamini-Hochberg procedure^75^, and the resulting q-values were reported alongside log2(fold changes) for each gene.

### CellTypist for Cell Type Prediction

CellTypist, a logistic regression-based cell type annotation tool, was used to transfer labels from atlases of fibroblasts and immune cells to our fibroblast and Treg datasets respectively^17^. First, we fit Celltypist to reference pre-annotated atlases using default parameters. To annotate our Treg datasets, a model was first fit to the CD4 T cell compartment of Hao et al., excluding CD4 proliferating cells^16^. In order to annotate fibroblasts according to tissue type, the model was fit to the entirety of Beuchler et al.^21^. Another final CellTypist model was fit to the entirety of Smillie et al. which was then used to characterize fibroblast subtype^22^. SGD-based feature selection was performed to fit models to Beuchler and Smillie atlases. Once fit, the models were used to predict individual cell types, and the predicted cell labels were recorded. Confidence scores for each prediction class are reported in extended data figures 1A and 2A.

### Pseudotime Ordering of Fibroblasts using Diffusion Maps

Diffusion pseudotime maps of time-course fibroblasts was performed using Pegasus as described in Haghverdi et al., 2016^76^. The affinity matrix was calculated using the top 50 PCs calculated for the time-course fibroblasts. Using this affinity matrix, 100 diffusion components were calculated, which were then used to generate single-cell pseudotime orderings. Fibroblasts from the 0-hour time point were used as the root for pseudotime calculation, and diffusion pseudotimes were normalized from 0 to 1.

### Transcription factor activity inference

Transcription factor activity scores were inferred using decoupler’s univariate linear model method^77^. The CollecTRI database was used to link transcription factors to downstream genes^78^.

### Analysis of PERTURB-LINK in BMDMs

Initial quality control of the Perturb-seq data was performed using empty CCEs to define filtering thresholds. A minimum feature count cutoff was established by calculating the mean plus 1.5 times the standard deviation of the feature counts within the empty CCE population. CCEs below this threshold were excluded. gRNA assignment was then conducted using MBC-based filtering, requiring a minimum of 3 MBCs per gRNA. Only cells containing a single gRNA were retained for downstream analysis. To integrate data across experimental batches, the Harmony algorithm^72^ was applied to the first 30 principal components (PCs) of the normalized transcriptomic data, using the experiment of origin as the batch variable. This corrected for technical artifacts while preserving biological variation.

### Cell Viability Analysis

To assess the effect of each perturbation on cell viability, we analyzed the relative abundance of cells within the LPS-stimulated condition. First, a gRNA-by-lane count matrix was generated by counting the number of cells assigned to each specific gRNA within each technical replicate (lane). These raw counts were then normalized. The counts were adjusted to correct for the uneven representation of gRNAs in the initial pooled plasmid library. The normalized counts for each targeting gRNA were compared to the distribution of counts from the non-targeting control gRNAs using a paired t-test across the technical replicates. This comparison yielded a p-value and a log2 fold change for each perturbation. Perturbations were classified as significantly cytotoxic or proliferative if the p-value was less than 0.05 and the absolute log2 fold change exceeded a 0.5 threshold.

### Generalized Linear Model and Regularization

To model the transcriptional effects of each perturbation, we used glmGamPoi^50^ Separate models were fitted for each gRNA, with cells containing non-targeting gRNAs serving as the reference control group. The experimental batch was included as a covariate in the model, and a likelihood ratio test was used to assess differential gene expression. The resulting beta coefficients, representing log2 fold changes, were regularized using an empirical Bayes shrinkage procedure. To achieve this, we applied a spike-and-slab prior, consisting of a point-mass spike and Laplace slab, to the LFCs After fitting the parameters of the spike-and-slab prior via maximum likelihood, we then computed the posterior mean of the LFCs, which served as the shrunken effect size for all downstream analyses. Finally, to explore the functional relationships between perturbations, a correlation analysis was performed on the shrunken beta matrix. PCA was performed on this scaled matrix, and the first 10 PCs were used to calculate the centered cosine similarity (Pearson correlation coefficient) between perturbations.

### Morphological perturbation analysis

Integration of morphology and perturbation metadata was performed at the CCE level. The first 50 PCs explained all the variance and all downstream morphology tests used this representation. Perturbation effects on morphology were quantified by comparing perturbed cells to matched NTCs using the energy distance statistic. Statistical significance was evaluated with a stratified permutation test, permuting labels within experiment–lane to control for batch/technical effects. For each perturbation we recorded the observed effect size and the permutation p-value. Unless otherwise specified, p-values are unadjusted; where reported, multiple testing was controlled with Benjamini–Hochberg at 5%.

### CRISPR–IF perturbation modeling (CD40 = green; PD-L1 = red)

Per-cell immunofluorescence intensities were analyzed using custom Python code. Within LPS + we fit a ordinary least squares model to each channel independently with categorical encoding for gRNA and experimental batch:

Q(outcome) ∽ C(gRNA) + C(experiment).

The non-targeting gRNA was set as the reference level to enable direct interpretation of coefficients as the effect of each perturbation relative to non-targeting, adjusted for experiment (batch). For every gRNA and channel, we report the log fold change, the two-sided Wald p-value, and the 95% Wald confidence interval.

## EXTENDED DATA

**EXTENDED DATA 1:**
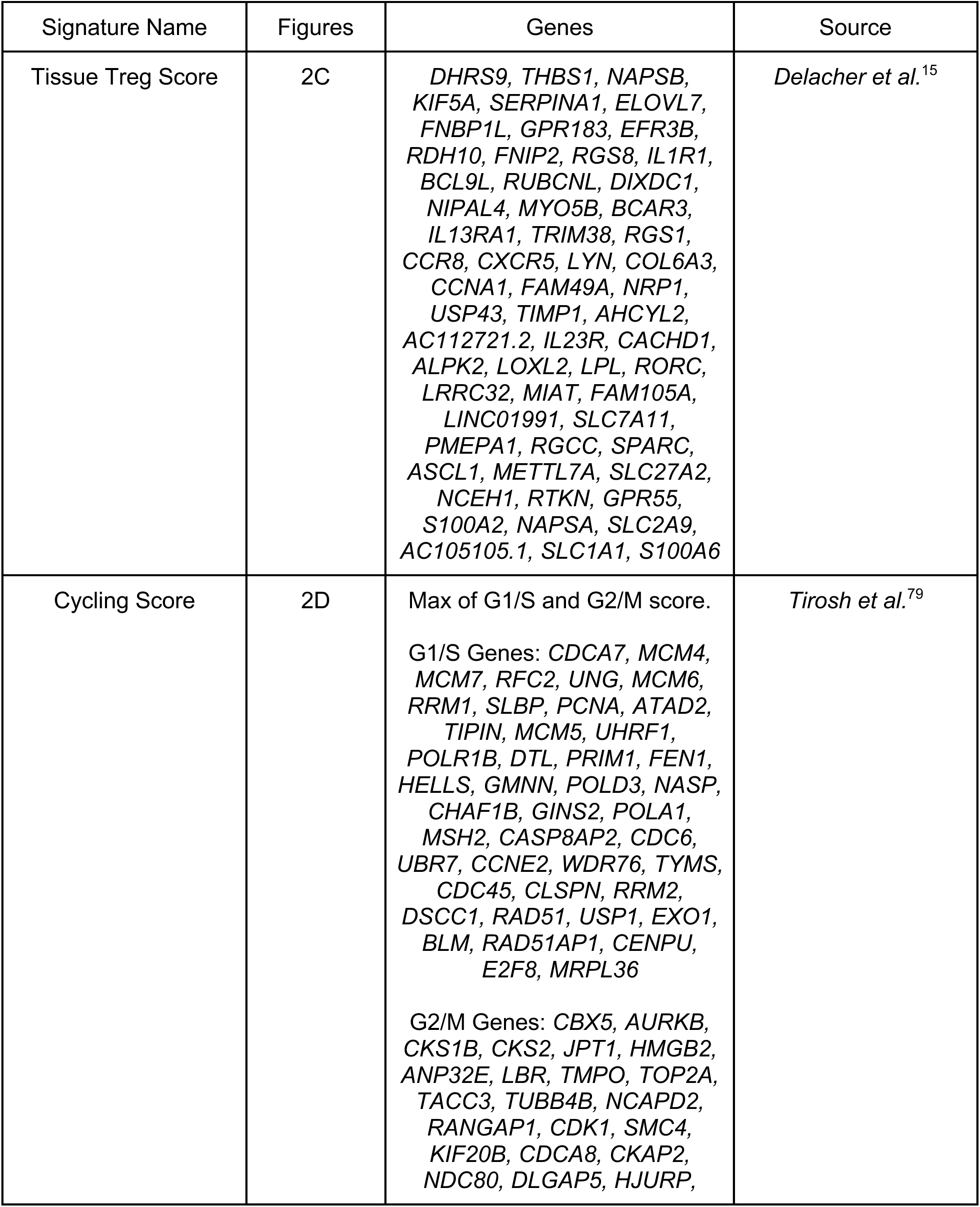
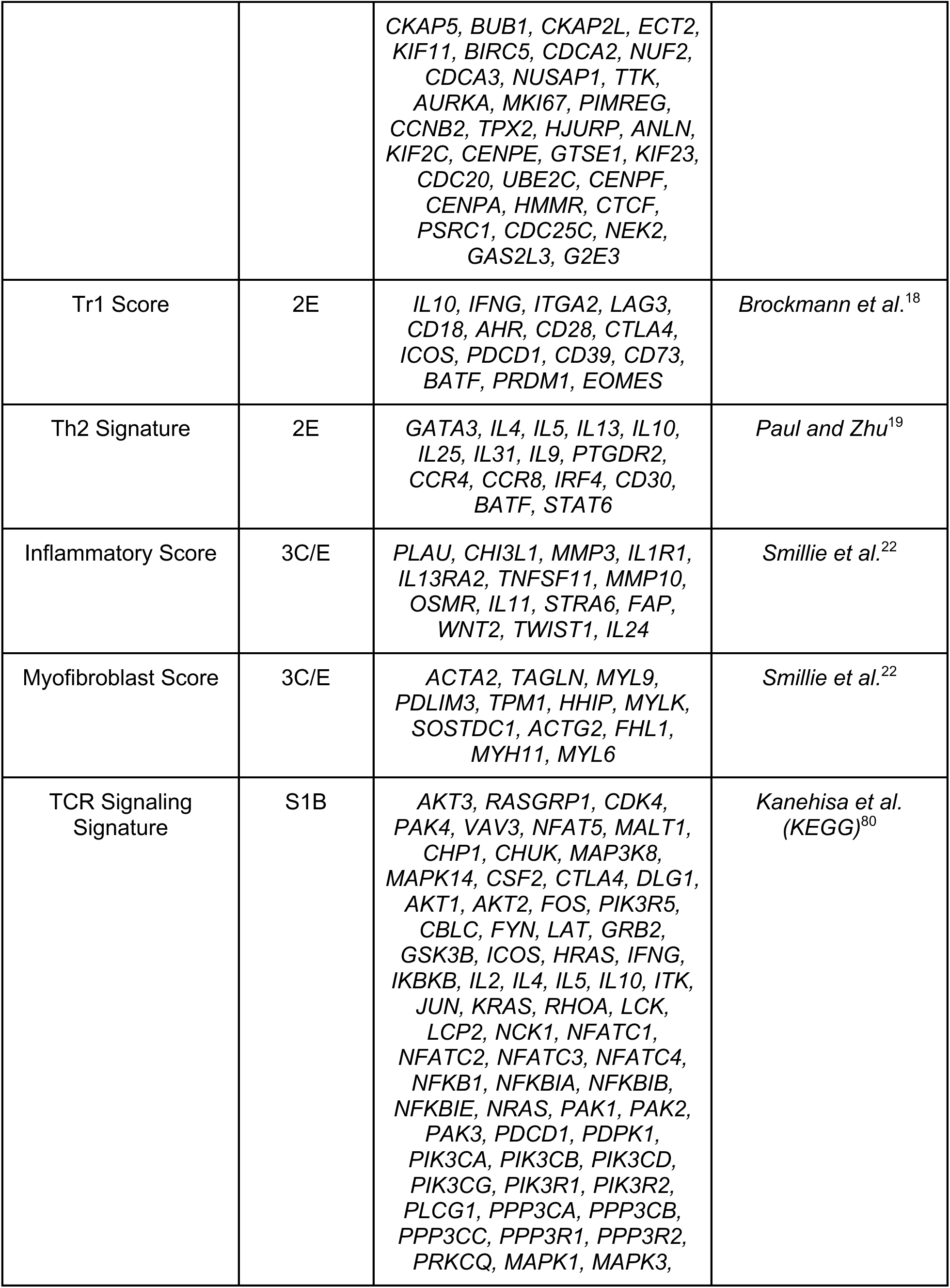
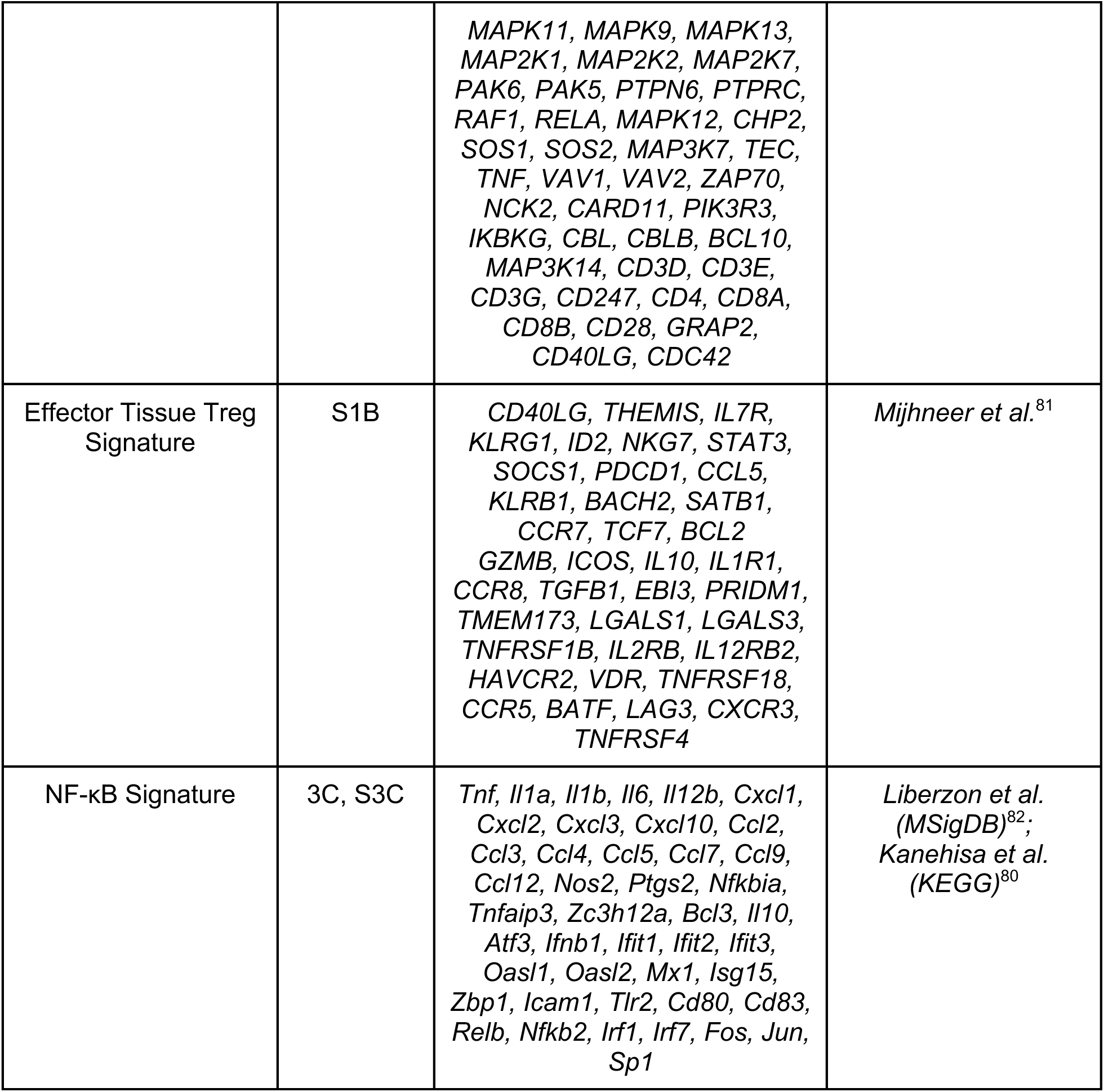
Gene signature gene lists and sources.

**Extended Data Fig. 1:**
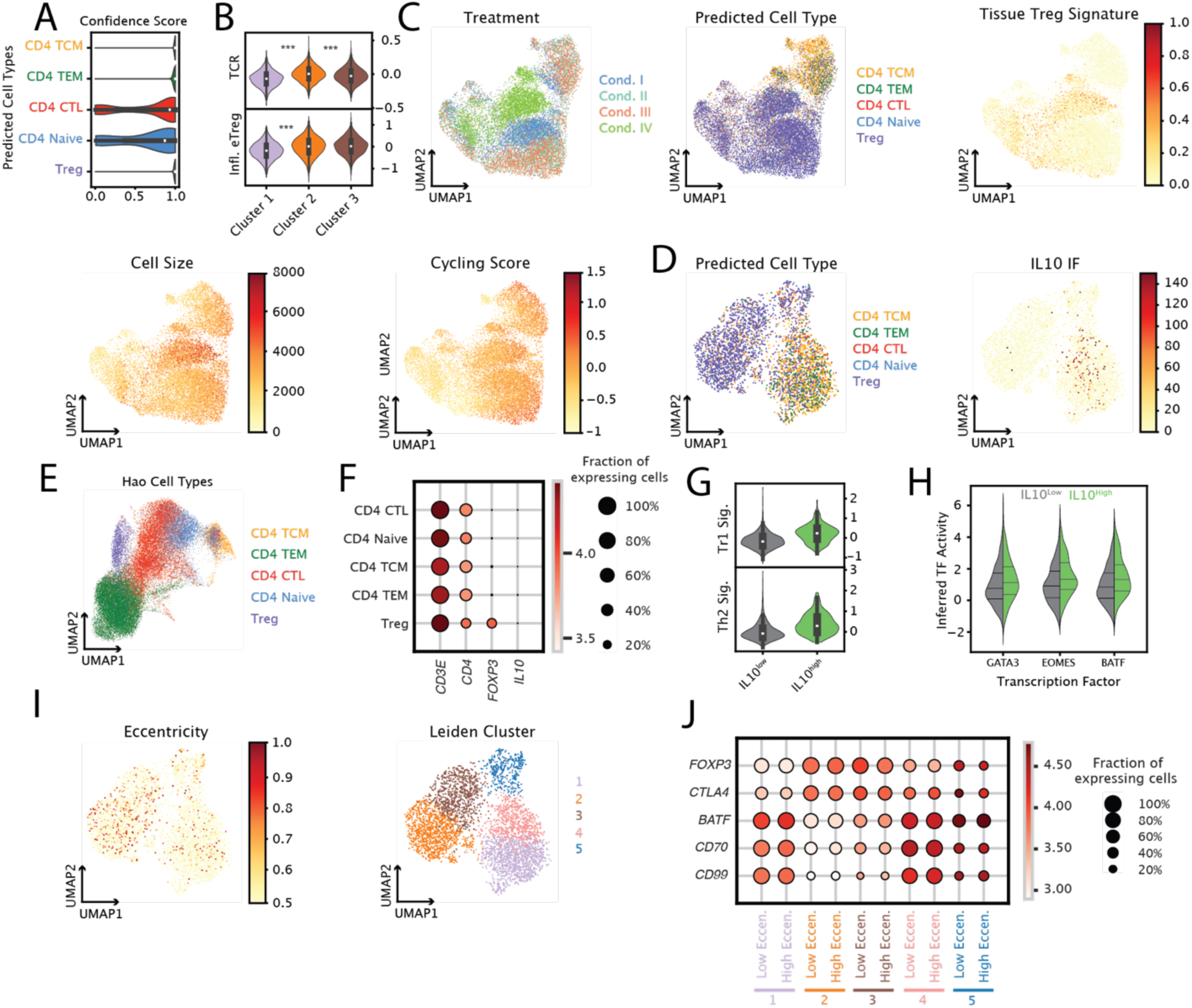
Joint proteomic, morphological, and transcriptomic profiling of human Tregs. **A**, Confidence scores of CellTypist predictions for each CD4 T class included in Hao et al. PBMC atlas^16^ (all cells included). **B**, T cell receptor (TCR), and inflammatory T reg signature scores for Leiden clusters 1-3 in Fig. 2B (see methods). **C**, UMAP of 17,967 T cells profiled at 6 days colored by (left to right, top to bottom) experimental condition, CellTypist predicted cell type, tissue Treg score, cell size, and cycling score. **D**, UMAP of 3,955 T cells profiled at 13 days colored by (left to right) CellTypist predicted cell type, and IL-10 IF signal. **E,** UMAP of Hao et al. CD4 cells colored by (left to right) cell type, and *IL10* expression. **F,** Dot plot of gene expression across cell types in Hao CD4 T cell atlas. **G**, Tr1 and Th2 signature scores for IL-10 IF high (IL-10 IF > 10) and IL-10 IF low (IL-10 IF < 10) T conventional cells (see methods). **H**, Inferred transcription factor activity score for IL-10 IF high and low Tconv cells. Transcription factor inference was performed using a univariate linear model from decoupler and the CollecTRI transcription factor database. **I**, UMAP of cells profiled at 13 days colored by (left to right) eccentricity, and Leiden cluster. **J,** Dot plot of gene expression for high (eccentricity > 0.5) and low eccentricity (eccentricity < 0.5) cells in each Leiden cluster in H. P values for gene score comparison and DE analysis were calculated using a two-sided Wilcoxon rank sum test. *\*Padj<0.05, **Padj<0.01, ***Padj<0.001*.

**Extended Data Fig. 2:**
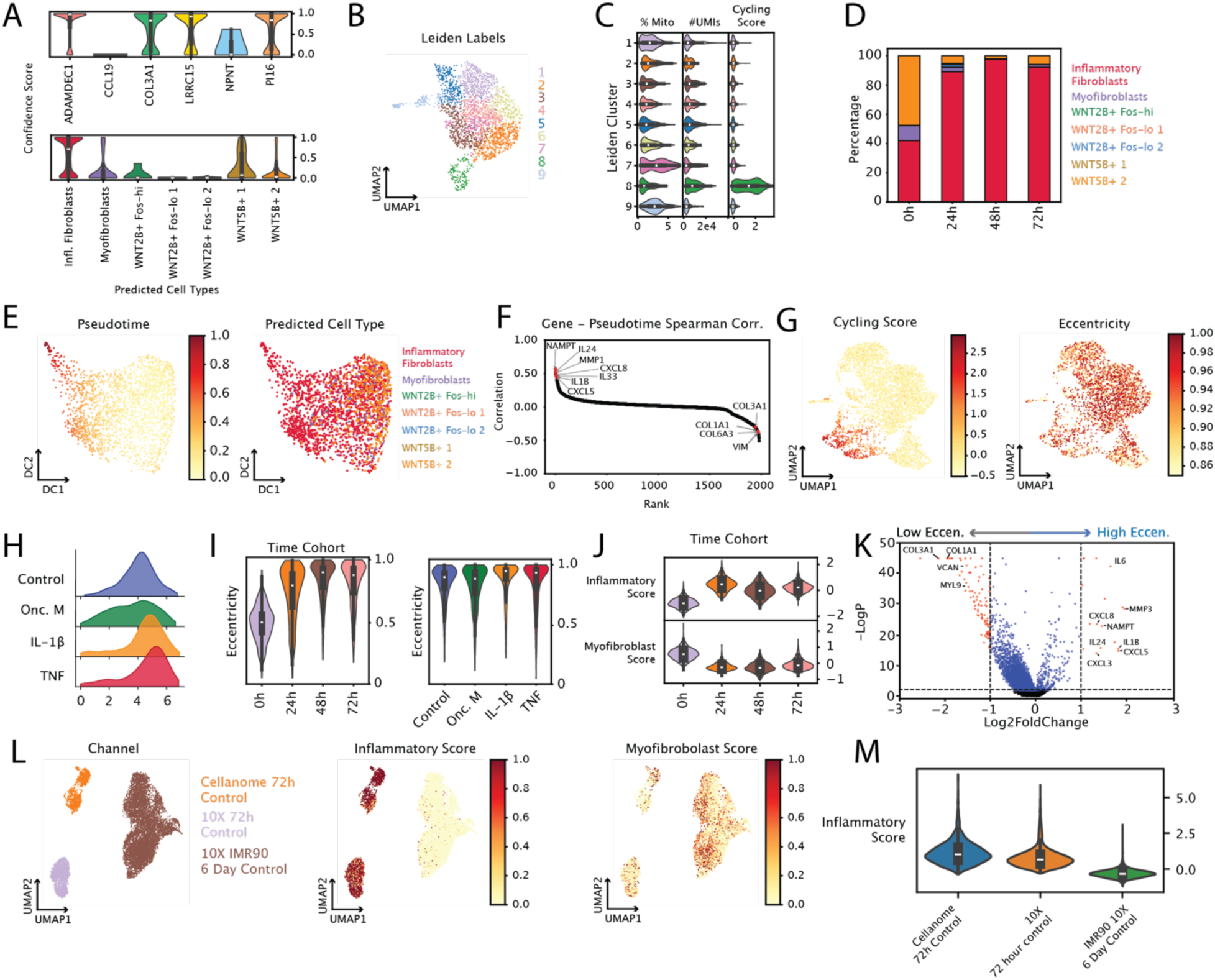
Joint proteomic, morphological, and transcriptomic profiling of human colonic fibroblasts. **A**, Confidence scores of CellTypist predictions for each fibroblast cell type included in Buechler et al.^21^ (top) and Smillie et al.^22^ (bottom) fibroblast atlases for all cells. **B**, UMAP of batch corrected transcriptomics of 2,559 time-course fibroblasts colored by Leiden cluster label. **C,** Percent mitochondrial RNA, number of MBCs, and cycling signature score for each cluster in B. **D**, Composition plot of the CellTypist-predicted cell types of fibroblasts profiled at each timepoint using the Smillie et al. fibroblast atlas. **E**, Diffusion map embedding of time course fibroblasts colored by (left to right) inferred pseudotime, and CellTypist predicted cell types from Smillie et al. atlas. **F**, Rank-ordered spearman correlation of spearman pseudotime and gene expression for time-course fibroblasts. **G,** UMAP of 4,598 fibroblasts, without batch correction, stimulated with various cytokines colored by (left to right) cycling signature score, and eccentricity. **H**, KDE plots of surface PDGFRa IF signal for cells treated with various cytokines. **I**, Violin plot of eccentricity for time course fibroblasts (left) and cytokine-treated fibroblasts (right). **J**, Inflammatory fibroblast and myofibroblast signature scores for time-course fibroblasts. **K**, Differential expression (DE) analysis between highly eccentric fibroblasts (eccentricity > 0.95), and less eccentric fibroblasts (eccentricity < 0.95) in the time-course experiment. Red points: genes expressed in highly eccentric fibroblasts (right, FDR < 0.05, log2FC > 1), or genes expressed more in more circular fibroblasts (left, FDR < 0.05, log2FC < -1). **L** UMAP of 1,249 control-group Hs.675 colonic fibroblasts profiled using CCE, 2,229 untreated human Hs.675 colonic fibroblasts profiled using 10X, and untreated, proliferating 8,443 IMR90 normal lung fibroblasts profiled using 10X from Palikyras et al.^23^ colored by (left to right) data source, inflammatory signature score, and myofibroblast signature score. **M**, Violin plot of inflammatory fibroblast score by data source of fibroblasts described in K. P values for both DE analysis and tissue Treg score comparison were calculated as described in methods.

**Extended Data Fig. 3:**
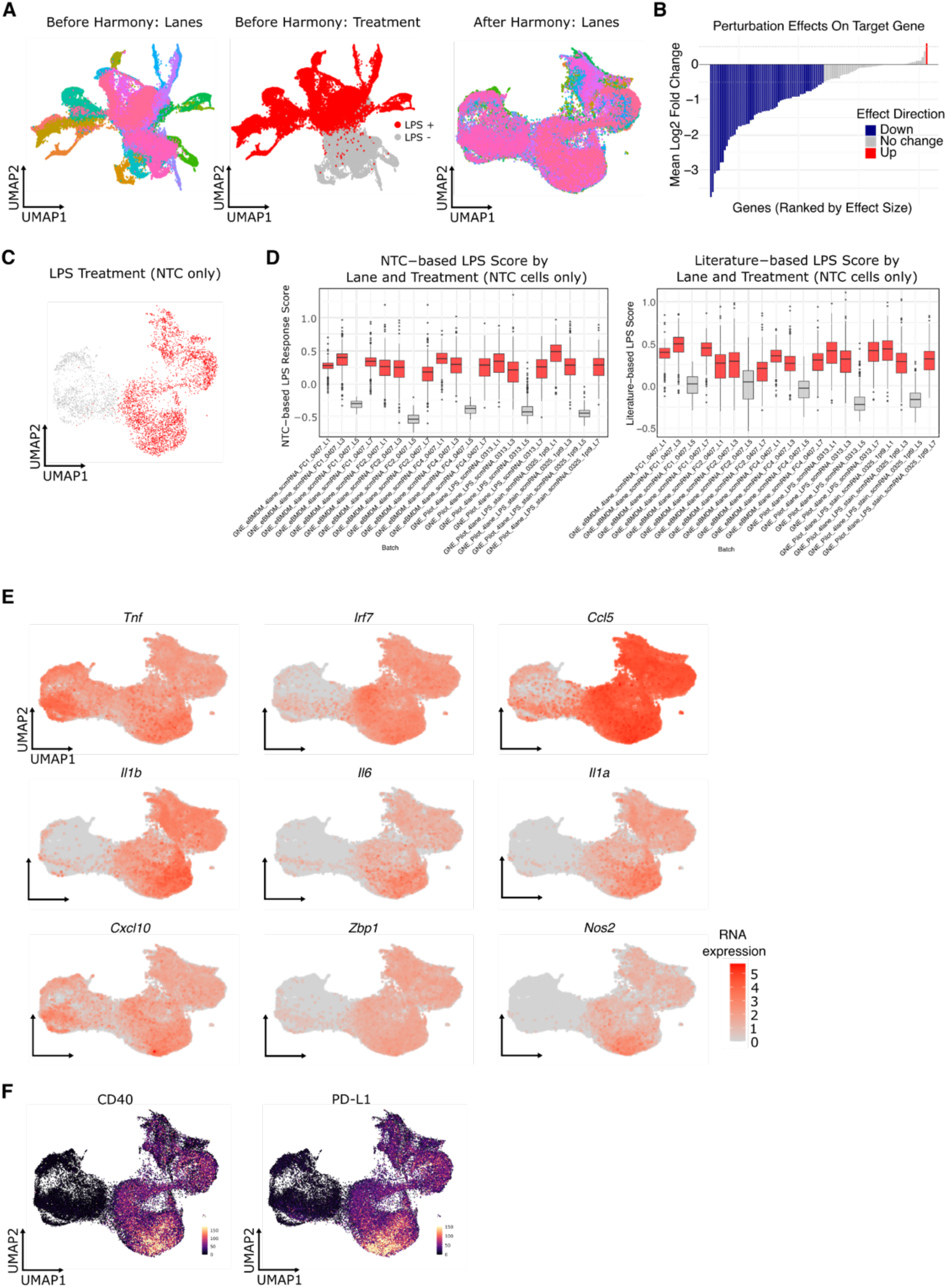
Batch correction, QC, and validation of multimodal data from the BMDM PERTURB-LINK screen. **A**, UMAPs before batch correction colored by experiment (left) and LPS status (middle), and a Harmony-corrected UMAP colored by experiment (right), showing removal of experiment-specific structure. **B,** Ranked Fold Change of Target Gene Expression vs. NTC. Blue (56) and Red (1) highlights indicate perturbations with the strongest changes in expression. **C**, Harmony-corrected UMAP restricted to non-targeting controls (NTC), colored by LPS status, confirming representation for signature scoring. **D**, Distributions of NTC-derived LPS response and literature-derived NF-κB signature scores, stratified by experiment and lane, demonstrating reproducibility across technical factors. **E**, Harmony-corrected UMAPs colored by expression of nine canonical inflammation genes. **F**, Harmony-corrected UMAPs with per-cell immunofluorescence (IF) intensity overlays for CD40 and PD-L1.

**Extended Data Fig. 4:**
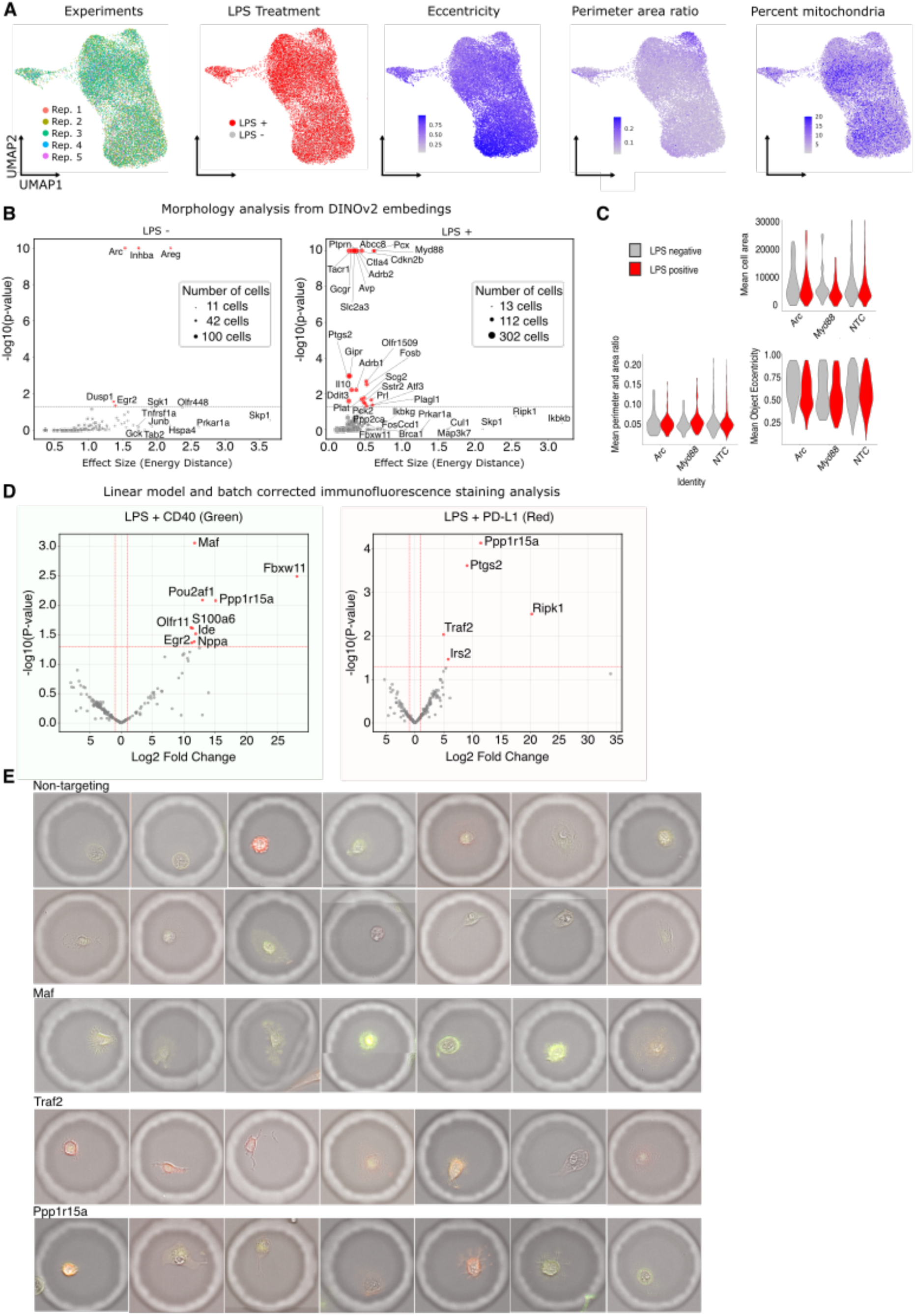
Morphology embeddings, effect sizes, and protein–morphology integration. **A**, UMAPs of DINOv2 morphology embeddings colored by experiment, LPS status, cell eccentricity, perimeter-to-area ratio, and percent mitochondria. The UMAP then has a clear trend of more circular cells in the bottom left while the top right describes cells that are long, narrow, or irregularly shaped. **B**, Morphology effect sizes computed from DINOv2 embeddings using energy distance for each perturbation versus non-targeting controls, shown by stimulus condition. Higher energy distance indicates greater deviation from control morphology. **C**, Violin plots of mean cell area, perimeter and area ratio, and eccentricity of Arc, Myd88 and NTC grouped by LPS stimuli. **D**, Linear-model–based, batch-corrected IF for LPS+ CD40 (green) and LPS+ PD-L1 (red), controlling for experiment; representative images and corresponding quantification are shown. **E**, Additional IF images for PD-L1 and CD40 hits identified by the linear model.

## REFERENCES

1. Yuan, J., Sheng, J. & Sims, P. A. SCOPE-Seq: a scalable technology for linking live cell imaging and single-cell RNA sequencing. Genome Biol. 19, 227 (2018).

2. Specht, H. et al. Single-cell proteomic and transcriptomic analysis of macrophage heterogeneity using SCoPE2. Genome Biol. 22, 50 (2021).

3. Bues, J. et al. Single-cell phenomics through integrated imaging and molecular profiling. 2025.11.28.690954 Preprint at 10.1101/2025.11.28.690954 (2025).

4. Xu, C. K. et al. Direct, high-throughput linking of single cell imaging and gene expression. 2025.09.01.672798 Preprint at 10.1101/2025.09.01.672798 (2025).

5. Baronas, D. et al. High-throughput single-cell omics using semipermeable capsules. Science 391, 1138–1145 (2026).

6. Mazelis, I., Sun, H., Kulkarni, A., Torre, T. L. & Klein, A. M. Multistep genomics on single cells and live cultures in subnanoliter capsules. Science 391, 1130–1137 (2026).

7. Khurana. Scalable multi-modal longitudinal and concurrent measurements of live cells in dynamic enclosures. Submitted (2025).

8. Miyara, M., Ito, Y. & Sakaguchi, S. TREG-cell therapies for autoimmune rheumatic diseases. Nat. Rev. Rheumatol. 10, 543–551 (2014).

9. Saini, N. & Neelapu, S. S. CAR Treg cells: prime suspects in therapeutic resistance. Nat. Med. 28, 1755–1756 (2022).

10. Liston, A., Pasciuto, E., Fitzgerald, D. C. & Yshii, L. Brain regulatory T cells. Nat. Rev. Immunol. 24, 326–337 (2024).

11. Wardell, C. M., Boardman, D. A. & Levings, M. K. Harnessing the biology of regulatory T cells to treat disease. Nat. Rev. Drug Discov. 24, 93–111 (2025).

12. Sakaguchi, S. et al. Regulatory T Cells and Human Disease. Annu. Rev. Immunol. 38, 541–566 (2020).

13. Chinen, T. et al. An essential role for the IL-2 receptor in Treg cell function. Nat. Immunol. 17, 1322–1333 (2016).

14. Fu, S. et al. TGF-β Induces Foxp3 + T-Regulatory Cells from CD4 + CD25 − Precursors. Am. J. Transplant. 4, 1614–1627 (2004).

15. Delacher, M. et al. Single-cell chromatin accessibility landscape identifies tissue repair program in human regulatory T cells. Immunity 54, 702–720.e17 (2021).

16. Hao, Y. et al. Integrated analysis of multimodal single-cell data. Cell 184, 3573–3587.e29 (2021).

17. Domínguez Conde, C., et al. Cross-tissue immune cell analysis reveals tissue-specific features in humans. Science 376, eabl5197 (2022).

18. Brockmann, L. et al. Molecular and functional heterogeneity of IL-10-producing CD4+ T cells. Nat. Commun. 9, 5457 (2018).

19. Paul, W. E. & Zhu, J. How are TH2-type immune responses initiated and amplified? Nat. Rev. Immunol. 10, 225–235 (2010).

20. West, N. R. et al. Oncostatin M drives intestinal inflammation and predicts response to tumor necrosis factor–neutralizing therapy in patients with inflammatory bowel disease. Nat. Med. 23, 579–589 (2017).

21. Buechler, M. B. et al. Cross-tissue organization of the fibroblast lineage. Nature 593, 575–579 (2021).

22. Smillie, C. S. et al. Intra- and Inter-cellular Rewiring of the Human Colon during Ulcerative Colitis. Cell 178, 714–730.e22 (2019).

23. Palikyras, S. et al. Rapid and synchronous chemical induction of replicative-like senescence via a small molecule inhibitor. Aging Cell 23, e14083 (2024).

24. Lawrence, T. & Natoli, G. Transcriptional regulation of macrophage polarization: enabling diversity with identity. Nat. Rev. Immunol. 11, 750–761 (2011).

25. Biswas, S. K. & Mantovani, A. Macrophage plasticity and interaction with lymphocyte subsets: cancer as a paradigm. Nat. Immunol. 11, 889–896 (2010).

26. Kudo, T. et al. Multiplexed, image-based pooled screens in primary cells and tissues with PerturbView. Nat. Biotechnol. 1–10 (2024) doi:10.1038/s41587-024-02391-0.

27. Zhang, Q., Lenardo, M. J. & Baltimore, D. 30 Years of NF-κB: A Blossoming of Relevance to Human Pathobiology. Cell 168, 37–57 (2017).

28. Lu, Y.-C., Yeh, W.-C. & Ohashi, P. S. LPS/TLR4 signal transduction pathway. Cytokine 42, 145–151 (2008).

29. Kawai, T. & Akira, S. Signaling to NF-κB by Toll-like receptors. Trends Mol. Med. 13, 460–469 (2007).

30. Pahl, H. L. Activators and target genes of Rel/NF-κB transcription factors. Oncogene 18, 6853–6866 (1999).

31. Hayden, M. S. & Ghosh, S. Shared Principles in NF-κB Signaling. Cell 132, 344–362 (2008).

32. Schambach, A. et al. Woodchuck hepatitis virus post-transcriptional regulatory element deleted from X protein and promoter sequences enhances retroviral vector titer and expression. Gene Ther. 13, 641–645 (2006).

33. Parnas, O. et al. A Genome-wide CRISPR Screen in Primary Immune Cells to Dissect Regulatory Networks. Cell 162, 675–686 (2015).

34. Deguine, J. & Barton, G. M. MyD88: a central player in innate immune signaling. F1000Prime Rep. 6, 97 (2014).

35. Shibuya, H. et al. TAB1: An Activator of the TAK1 MAPKKK in TGF-β Signal Transduction. Science 272, 1179–1182 (1996).

36. Sakurai, H. Targeting of TAK1 in inflammatory disorders and cancer. Trends Pharmacol. Sci. 33, 522–530 (2012).

37. Winston, J. T. et al. The SCFβ-TRCP–ubiquitin ligase complex associates specifically with phosphorylated destruction motifs in IκBα and β-catenin and stimulates IκBα ubiquitination in vitro. Genes Dev. 13, 270–283 (1999).

38. Liu, T., Zhang, L., Joo, D. & Sun, S.-C. NF-κB signaling in inflammation. Signal Transduct. Target. Ther. 2, 17023 (2017).

39. Hinz, M. & Scheidereit, C. The IκB kinase complex in NF-κB regulation and beyond. EMBO Rep. 15, 46–61 (2014).

40. Etemadi, N., et al. TRAF2 regulates TNF and NF-κB signalling to suppress apoptosis and skin inflammation independently of Sphingosine kinase 1. eLife 4, e10592 (2015).

41. Dixit, A. et al. Perturb-Seq: Dissecting Molecular Circuits with Scalable Single-Cell RNA Profiling of Pooled Genetic Screens. Cell 167, 1853–1866.e17 (2016).

42. Oquab, M., et al. DINOv2: Learning Robust Visual Features without Supervision. Preprint at 10.48550/arXiv.2304.07193 (2024).

43. Peidli, S. et al. scPerturb: harmonized single-cell perturbation data. Nat. Methods 21, 531–540 (2024).

44. Ufer, F., et al. Arc/Arg3.1 governs inflammatory dendritic cell migration from the skin and thereby controls T cell activation. Sci. Immunol. 1, eaaf8665–eaaf8665 (2016).

45. Tintelnot, J. et al. Arc/Arg3.1 defines dendritic cells and Langerhans cells with superior migratory ability independent of phenotype and ontogeny in mice. Eur. J. Immunol. 49, 724–736 (2019).

46. Xie, P. TRAF molecules in cell signaling and in human diseases. J. Mol. Signal. 8, 7 (2013).

47. Bishop, G. A. The multifaceted roles of TRAFs in the regulation of B-cell function. Nat. Rev. Immunol. 4, 775–786 (2004).

48. Clavarino, G. et al. Protein phosphatase 1 subunit Ppp1r15a/GADD34 regulates cytokine production in polyinosinic:polycytidylic acid-stimulated dendritic cells. Proc. Natl. Acad. Sci. 109, 3006–3011 (2012).

49. Marciniak, S. J. et al. CHOP induces death by promoting protein synthesis and oxidation in the stressed endoplasmic reticulum. Genes Dev. 18, 3066–3077 (2004).

50. Ahlmann-Eltze, C. & Huber, W. glmGamPoi: fitting Gamma-Poisson generalized linear models on single cell count data. Bioinformatics 36, 5701–5702 (2021).

51. Denisenko, E. et al. Systematic assessment of tissue dissociation and storage biases in single-cell and single-nucleus RNA-seq workflows. Genome Biol. 21, 130 (2020).

52. Shalek, A. K. et al. Single-cell RNA-seq reveals dynamic paracrine control of cellular variation. Nature 510, 363–369 (2014).

53. Janesick, A. et al. High resolution mapping of the tumor microenvironment using integrated single-cell, spatial and in situ analysis. Nat. Commun. 14, 8353 (2023).

54. Lopez, T. et al. High-Throughput Multiomics Profiling of Model Systems Using the AVITI24 Platform. 2025.05.03.651997 Preprint at 10.1101/2025.05.03.651997 (2025).

55. Merritt, C. R. et al. Multiplex digital spatial profiling of proteins and RNA in fixed tissue. Nat. Biotechnol. 38, 586–599 (2020).

56. Pitino, E. et al. STAMP: Single-cell transcriptomics analysis and multimodal profiling through imaging. Cell 0, (2025).

57. Feldman, D. et al. Optical Pooled Screens in Human Cells. Cell 179, 787–799.e17 (2019).

58. Chen, W. et al. Live-seq enables temporal transcriptomic recording of single cells. Nature 608, 733–740 (2022).

59. Marcuccio, F. et al. Single-cell nanobiopsy enables multigenerational longitudinal transcriptomics of cancer cells. Sci. Adv. 10, eadl0515 (2024).

60. Liao, S. et al. Stereo-cell: Spatial enhanced-resolution single-cell sequencing with high-density DNA nanoball-patterned arrays. Science 10.1126/science.adr0475 (2025) doi:10.1126/science.adr0475.

61. Li, H. et al. A comprehensive benchmarking with practical guidelines for cellular deconvolution of spatial transcriptomics. Nat. Commun. 14, 1548 (2023).

62. Cable, D. M. et al. Robust decomposition of cell type mixtures in spatial transcriptomics. Nat. Biotechnol. 40, 517–526 (2022).

63. Biancalani, T. et al. Deep learning and alignment of spatially resolved single-cell transcriptomes with Tangram. Nat. Methods 18, 1352–1362 (2021).

64. Argelaguet, R. et al. MOFA+: a statistical framework for comprehensive integration of multi-modal single-cell data. Genome Biol. 21, 111 (2020).

65. Ashuach, T. et al. MultiVI: deep generative model for the integration of multimodal data. Nat. Methods 20, 1222–1231 (2023).

66. González, I., Déjean, S., Martin, P. G. P. & Baccini, A. CCA: An R Package to Extend Canonical Correlation Analysis. J. Stat. Softw. 23, 1–14 (2008).

67. Kaminow, B., Yunusov, D. & Dobin, A. STARsolo: accurate, fast and versatile mapping/quantification of single-cell and single-nucleus RNA-seq data. 2021.05.05.442755 Preprint at 10.1101/2021.05.05.442755 (2021).

68. Redmon, J., Divvala, S., Girshick, R. & Farhadi, A. You Only Look Once: Unified, Real-Time Object Detection. Preprint at 10.48550/arXiv.1506.02640 (2016).

69. He, K., Gkioxari, G., Dollár, P. & Girshick, R. Mask R-CNN. Preprint at 10.48550/arXiv.1703.06870 (2018).

70. Li, B. et al. Cumulus provides cloud-based data analysis for large-scale single-cell and single-nucleus RNA-seq. Nat. Methods 17, 793–798 (2020).

71. lilab-bcb/harmony-pytorch. lilab-bcb (2025).

72. Korsunsky, I. et al. Fast, sensitive and accurate integration of single-cell data with Harmony. Nat. Methods 16, 1289–1296 (2019).

73. McInnes, L., Healy, J. & Melville, J. UMAP: Uniform Manifold Approximation and Projection for Dimension Reduction. Preprint at 10.48550/arXiv.1802.03426 (2020).

74. Traag, V. A., Waltman, L. & van Eck, N. J. From Louvain to Leiden: guaranteeing well-connected communities. Sci. Rep. 9, 5233 (2019).

75. Benjamini, Y. & Hochberg, Y. On the Adaptive Control of the False Discovery Rate in Multiple Testing With Independent Statistics. J. Educ. Behav. Stat. 25, 60–83 (2000).

76. Haghverdi, L., Büttner, M., Wolf, F. A., Buettner, F. & Theis, F. J. Diffusion pseudotime robustly reconstructs lineage branching. Nat. Methods 13, 845–848 (2016).

77. Badia-i-Mompel, P., et al. decoupleR: ensemble of computational methods to infer biological activities from omics data. Bioinforma. Adv. 2, vbac016 (2022).

78. Müller-Dott, S. et al. Expanding the coverage of regulons from high-confidence prior knowledge for accurate estimation of transcription factor activities. Nucleic Acids Res. 51, 10934–10949 (2023).

79. Tirosh, I. et al. Single-cell RNA-seq supports a developmental hierarchy in human oligodendroglioma. Nature 539, 309–313 (2016).

80. Kanehisa, M., Furumichi, M., Sato, Y., Matsuura, Y. & Ishiguro-Watanabe, M. KEGG: biological systems database as a model of the real world. Nucleic Acids Res. 53, D672–D677 (2025).

81. Mijnheer, G. et al. Conserved human effector Treg cell transcriptomic and epigenetic signature in arthritic joint inflammation. Nat. Commun. 12, 2710 (2021).

82. Liberzon, A. et al. The Molecular Signatures Database Hallmark Gene Set Collection. Cell Syst. 1, 417–425 (2015).

